# Effector loss drives adaptation of Pseudomonas syringae pv. actinidiae to Actinidia arguta

**DOI:** 10.1101/2021.11.15.468702

**Authors:** Lauren M. Hemara, Jay Jayaraman, Paul W. Sutherland, Mirco Montefiori, Saadiah Arshed, Abhishek Chatterjee, Ronan Chen, Mark Andersen, Carl H. Mesarich, Otto van der Linden, Magan M. Schipper, Joel L. Vanneste, Cyril Brendolise, Matthew D. Templeton

## Abstract

A pandemic isolate of *Pseudomonas syringae* pv. *actinidiae* biovar 3 (Psa3) has devastated kiwifruit orchards growing cultivars of *Actinidia chinensis*. In contrast, *A. arguta* (kiwiberry) is resistant to Psa3. This resistance is mediated via effector-triggered immunity, as demonstrated by induction of the hypersensitive response in infected *A. arguta* leaves, observed by microscopy and quantified by ion-leakage assays. Isolates of Psa3 that cause disease in *A. arguta* have been isolated and analyzed, revealing a 49 kb deletion in the exchangeable effector locus (EEL). This natural EEL-mutant isolate and strains with synthetic knockouts of the EEL were more virulent in *A. arguta* plantlets than wild-type Psa3. Screening of a complete library of Psa3 effector knockout strains identified increased growth *in planta* for knockouts of four effectors – AvrRpm1a, HopF1c, HopZ5a, and the EEL effector HopAW1a – suggesting a resistance response in *A. arguta*. Hypersensitive response (HR) assays indicate that three of these effectors trigger a host species-specific HR. A Psa3 strain with all four effectors knocked out escaped host recognition, but a cumulative increase in bacterial pathogenicity and virulence was not observed. These avirulence effectors can be used in turn to identify the first cognate resistance genes in *Actinidia* for breeding durable resistance into future kiwifruit cultivars.

## Introduction

The *Pseudomonas syringae* species complex contains over 60 pathovars, each with a discrete host range (Bull et al., 2010; Morris et al., 2019; Xin et al., 2018). The collective host breadth of the *P. syringae* species complex makes this bacterial plant pathogen an ideal model for studying the molecular basis of host specificity. *P. syringae* pv. *actinidiae* (Psa), the causal agent of kiwifruit canker, is a recently emerged plant pathogen. The disease was first isolated from *A. chinensis* var. *deliciosa* (green-fleshed kiwifruit), and *A. arguta* (kiwiberry) in Japan in 1984 (Serizawa et al., 1989; Takikawa et al., 1989). There was a subsequent outbreak in South Korea in the mid 1990s (Koh et al., 1994). However, it was the emergence of a pandemic strain that spread rapidly around the world from 2008, which particularly devastated orchards of *Actinidia chinensis* var. *chinensis* (gold-fleshed kiwifruit) (Scortichini, 1994; Vanneste, 2017). Isolates from these three separate outbreaks of bacterial canker have been grouped into biovars and recently two more biovars have been described (Fujikawa and Sawada, 2016; Fujikawa and Sawada, 2019). Biovars of Psa have closely related core genes and are primarily distinguished by their variable accessory genomes, which include effectors and toxin biosynthesis clusters (Sawada and Fujikawa, 2019).

Psa3 is the only biovar present in New Zealand, alongside the closely related pathovar *P. syringae* pv. *actinidifoliorum* (Cunty et al., 2015). Psa was first detected in New Zealand’s kiwifruit-growing region of Te Puke in 2010 (Everett et al., 2011). The introduction of Psa3 to New Zealand orchards in 2010 appears to have been a single event, as the Psa population has remained largely clonal (Colombi et al., 2017). Subsequently, in response to copper application, acquisition of copper-resistance integrative conjugative elements, plasmids and transposons has been observed (Colombi et al., 2017).

Resistance to Psa3 has been observed within an *Actinidia* germplasm collection, including in *A. arguta* (Datson et al., 2013). In contrast, in a commercial *A. arguta* orchard, rare Psa infections of *A. arguta* ‘HortGem Tahi’ and ‘HortGem Rua’ cultivars produced symptomatic angular necrotic leaf spots; however, the outbreak did not result in a significant loss of orchard productivity (Vanneste et al., 2014). Additionally, limited infection is observed in *A. arguta* seedlings stab-inoculated with Psa, with infection limited to the tissue immediately surrounding the inoculation site (Hoyte et al., 2013). This appears to be related to earlier recognition of Psa3 in *A. arguta* than in *A. chinensis* (Nunes da Silva et al., 2020), suggesting that *A. arguta* has a degree of resistance to Psa, which may be conferred by undiscovered resistance genes recognizing Psa3 effectors.

Host range in the *P. syringae* species complex is largely driven by the composition of the effector complement, which consists of at least 68 effector families (Dillon et al., 2019). Effectors are thus intrinsic to the ability of specialized pathogens within this species complex to cause disease *in planta*. However, an individual *P. syringae* strain carries only a fraction of this “super pan effector repertoire”; further still, only a subset of these effectors, owing to redundancy within the effector complement, may make an indispensable contribution to virulence in a given host (Wei et al., 2015; Wei and Collmer, 2018).

Effector proteins are translocated into host cells via a type III secretion system (T3SS) encoded by the *hrp/hrc* gene cluster (Alfano et al., 2000). The *hrp/hrc* genes are required for the production of the T3SS, as Δ*hrcC* deletion mutants cannot deliver effector proteins into host plant cells, thus preventing pathogenicity in host plants (Alfano et al., 2000; Jayaraman et al., 2020). Once in host cells, effectors promote bacterial virulence by interacting with host targets to suppress host immunity, allowing the pathogen to invade host tissue, acquire nutrients and cause disease (Bent and Mackey, 2007; Chisholm et al., 2006; Zipfel, 2009). Plant resistance proteins monitor the integrity of, for example, defence signaling cascades, and can detect subversion by bacterial effectors, inducing effector-triggered immunity (ETI), thus restoring plant resistance (Chisholm et al., 2006; Ngou et al., 2021; Yuan et al., 2021). *P. syringae* T3S effectors are termed Hop (<u>H</u>rp <u>o</u>uter <u>p</u>rotein) or Avr (avirulence) proteins (Lindeberg et al., 2012). Avr proteins are a subset of Hop effectors that are recognized by the products of known plant disease resistance genes.

In the majority of *P. syringae* genomes, two groups of effectors are co-located with the *hrp/hrc* gene cluster, forming a tripartite pathogenicity island. These are the conserved effector locus (CEL) and the more variable exchangeable effector locus (EEL) (Alfano et al., 2000). CEL effectors are required for pathogenesis, demonstrated by strongly reduced pathogenicity and virulence in *P. syringae* ΔCEL strains in host plants (Alfano et al., 2000; Badel et al., 2006; Jayaraman et al., 2020). The EEL has been remodeled extensively between different *P. syringae* pathovars, creating significant genetic variation through mutation, insertion, deletion, and recombination (Xin et al., 2018). The non-syntenic EEL from Psa3 ICMP 18884 contains the effectors *hopQ1a*, *hopD1a*, *avrD1*, *avrB2b*, *hopAB1b*, *hopF4a*, *hopAW1a*, *hopF1e*, *hopAF1b*, *hopD2a*, and *hopF1a*. However, even within the Psa pathovar, the EEL is variable across biovars and strains (McCann et al., 2013; Rikkerink et al., 2015).

While previous research has identified that Psa3 CEL (and related) effectors are required for virulence (Jayaraman et al., 2020), no specific Psa3 avirulence effectors recognized by *Actinidia* spp. have been identified. In this study, a genome sequencing field survey and subsequent effector knockout assays identified four Psa3 avirulence effectors associated with the resistance response to Psa3 in *A. arguta* AA07_03: HopAW1a, AvrRpm1a, HopF1c, and HopZ5a.

## Results

### Psa3 induces the hypersensitive response in *Actinidia arguta*

Previous work (Jayaraman et al., 2021) showed that *A. arguta* plants were resistant to Psa3, associated with a quantifiable increase in ion leakage due to membrane disruption of the dying cells indicative of a hypersensitive response (HR). Leaves of *A. arguta* and *A. chinensis* var. *chinensis* spray-infected with Psa3 were visually inspected for macroscopic symptoms at 1 day post inoculation (dpi) (*A. arguta*) or 5 dpi (*A. chinensis* var. *chinensis*). This revealed small dark brown patches each consisting of a few cells, indicative of an HR in *A. arguta*, in contrast to leaves of *A. chinensis* var. *chinensis* (Figure 1A). Accumulated phenolic compounds, characteristic of the HR, were more obvious when the leaves were cleared (Figure 1A). At higher magnification, immuno-labelling with an antibody specific for β1-3-glucan revealed callose accumulation in the portion of the cell walls of live cells in direct contact with the dead cells in *A. arguta*, and a lack of cell death, but some callose deposition in *A. chinensis* var. *chinensis* (Figure 1B). Under a fluorescence microscope the dead mesophyll cells were readily visible because of their high concentrations of phenolic compounds following cell wall degradation. Collectively these results in *A. arguta* treated with Psa3 show hypersensitive cell death and a defence response in adjacent cells, hallmarks of ETI.

**Figure 1:**
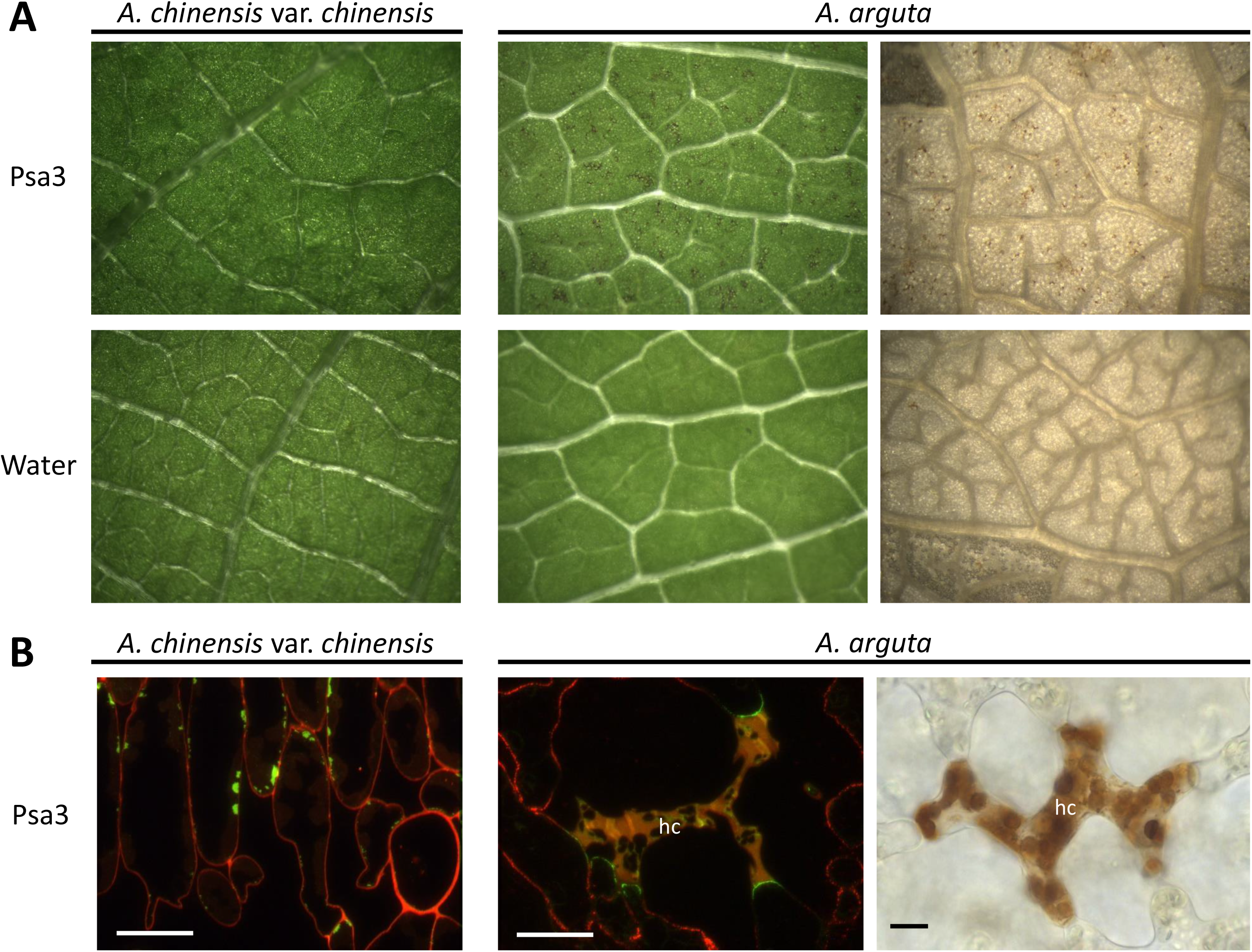
*Pseudomonas syringae* pv. *actinidiae* biovar3 (Psa3) induces the hypersensitive response in *Actinidia arguta.* *A. arguta* and *A. chinensis* var. *chinensis* leaves displaying symptoms following infection with Psa3 V-13 at 1 day post-infection (dpi) in *A. arguta* and 5 dpi in *A. chinensis* var. *chinensis*. **(A)** Visualization of macroscopically visible localized cell death indicative of a hypersensitive response (HR) to in leaves of *A. arguta*, in contrast to *A. chinensis* var. *chinensis*, spray-infected with Psa3 V-13 at 10^8^ cfu/mL or water control (left). *A. arguta* leaves were cleared in acetic acid:ethanol to better visualize brown phenolic compounds indicating cell death (right; brown speckling in the images). **(B)** Fluorescence microscopy of Psa3 V-13-infected *A. arguta* and *A. chinensis* var. *chinensis* mesophyll tissue. Callose (β1-3-glucan) is immuno-labelled and fluorescence indicated in green; cell wall pectin is immuno-labeled and fluorescence indicated in red; yellow coloring is accumulation of phenolic compounds in cells showing hypersensitive cell death (hc; left) and loss of cell wall integrity. Bright field microscopy of cleared *A. arguta* leaf in **A** indicates phenolic compound accumulation in cells showing hypersensitive cell death (hc; right) caused by cell wall breakdown. Scale bars represent 10 µm.

### Psa isolates from symptomatic *A. arguta* leaves have a 49 kb deletion in the exchangeable effector locus

During routine surveys of our *Actinidia* spp. germplasm collection at the Te Puke Research Orchard, we observed leaves of *A. arguta* ‘HortGem Tahi’ with leaf spot disease symptoms. The leaf spots comprised an angular necrotic zone surrounded by a chlorotic halo (Figure 2A). *P. syringae* was isolated from these lesions and confirmed to be Psa3 using qPCR (Andersen et al., 2017; Barrett-Manako et al., 2021). Several of these isolates were sequenced using the Illumina HiSeq platform. Four isolates had a 49 kb deletion in the EEL, with the deletion flanked by miniature inverted-repeat transposable elements (MITEs). The deleted region contained several effectors including *hopAW1a, hopF1e, hopAF1b, hopD2a,* and *hopF1a*, and genes encoding a putative novel non-ribosomal peptide synthase (NRPS) toxin synthesis pathway (Figure 2B). One of the isolates with the 49 kb deletion, Psa3 X-27, was checked by PCR spanning the deletion site (*Psa-X27*; effector check-primers Table S1) and Sanger sequencing to confirm the deletion (Figure 2C).

**Figure 2:**
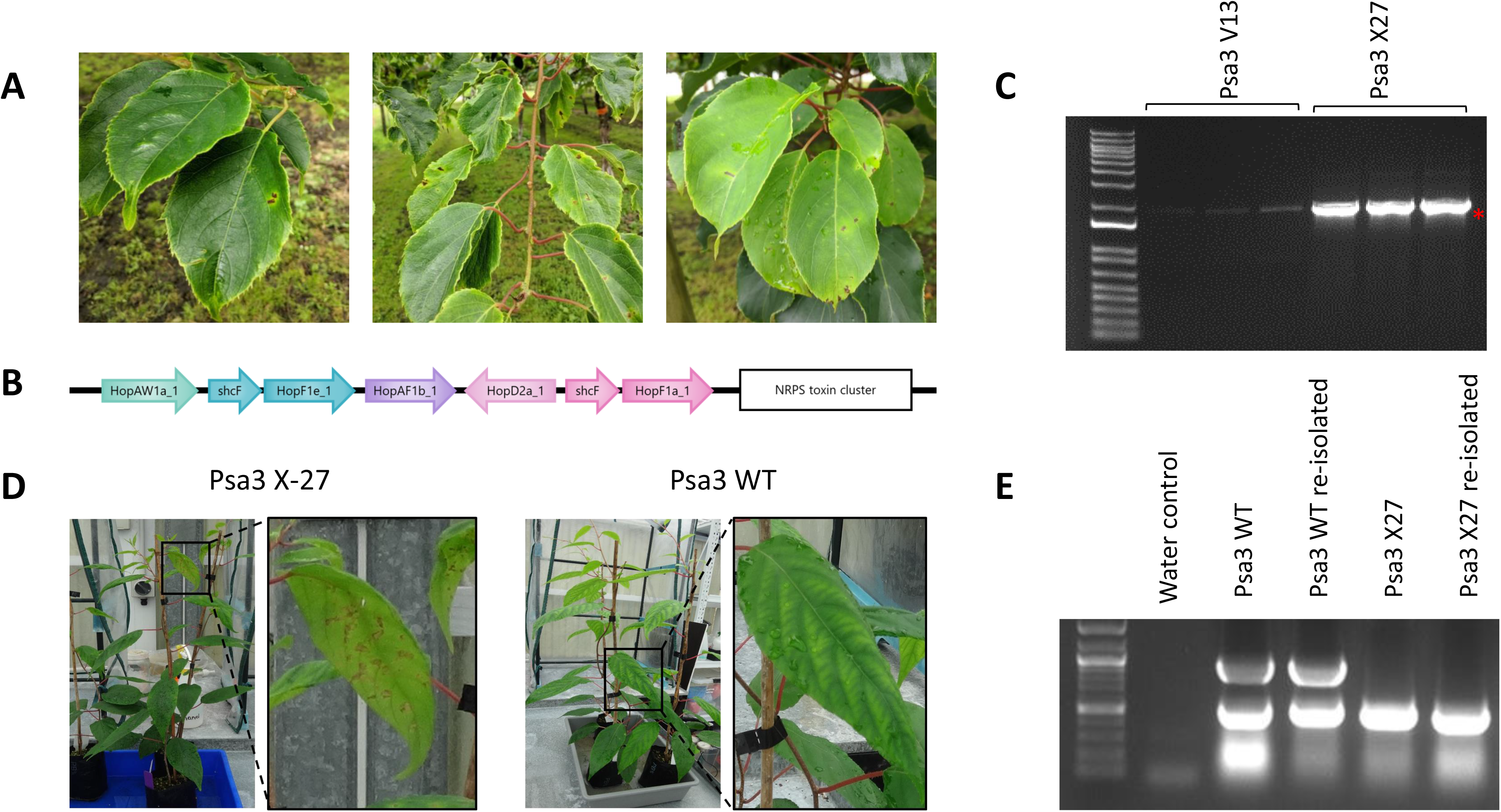
*Pseudomonas syringae* pv. *actinidiae* biovar3 (Psa3) isolated from symptomatic *Actinidia arguta* plants has a deletion in the exchangeable effector locus. **(A)** Psa leaf spot symptoms on commercial *A. arguta* ‘HortGem Tahi’ plants in the Plant & Food Research Te Puke Research Orchard. **(B)** The Psa3 X-27 gene deletion spans the effectors *hopAW1a, hopF1e, hopAF1b, hopD2a, hopF1a*, and the non-ribosomal peptide synthase (NRPS) toxin cluster. The Psa3 X-27 gene deletion was identified through whole-genome sequencing on an Illumina HiSeq platform and confirmed by PCR. **(C)** Three colonies of Psa3 ICMP 18884 (V13) or Psa3 X-27 were used as templates for PCR across the deletion boundary *Psa-X27* (1804 bp) and the band indicating deletion (red asterisk) confirmed by Sanger sequencing. DNA marker is 1Kb Plus DNA Ladder from Thermo Fisher (NZ). **(D)** Psa3 X-27 or Psa3 ICMP 10627 (WT) were sprayed onto potted *A. arguta* AA07_03 plants and photographs of symptoms taken 6 months post-infection. **(E)** Psa3 ICMP 10627 (WT) and Psa3 X-27 re-isolated from infected leaves and confirmed by multiplex PCR for *Psa-ompP1* (492 bp) and the EEL effector gene *hopF1e* (883 bp). DNA ladder is 100bp DNA Marker™ from Zymo Research (USA).

Potted plants of *A. arguta* AA07_03 were infected with Psa3 ICMP 10627 (wild type, WT) and one of the isolates with the 49 kb deletion, Psa3 X-27. Leaves of these Psa3 X-27-infected plants had chlorotic halos and necrotic leaf symptoms, in contrast to plants infected with Psa3 (WT), which displayed no visible symptoms (Figure 2D). Psa3 X-27 and Psa3 WT were re-isolated from the spray-infected leaves and verified by PCR. Here, confirmation was achieved by multiplex PCR for EEL locus effector gene *hopF1e* (883 bp) and Psa-*ompP1* primers (Koh and Nou, 2002) (492 bp), with both present in the WT but only Psa-*ompP1* present in the original and the re-isolated X-27 (Figure 2E).

### Psa3 X-27 escapes host recognition in *A. arguta* through effector loss

To determine whether it was the Psa3 X-27 multi-effector deletion that allowed this isolate to overcome *A. arguta* resistance, the Psa3 V-13 Δ*sEEL* knockout strain was developed to have the same EEL effector deletion as Psa3 X-27 while retaining the putative NRPS toxin biosynthesis gene cluster. *A. arguta* AA07_03 plantlets were flood-inoculated with Psa3 V-13, Psa3 X-27, and Psa3 V-13 Δ*sEEL* and assessed for *in planta* growth for a single experimental run (Figure 3). Infected plantlets were sampled at 6 and 12 dpi. Psa3 V-13 triggered resistance in *A. arguta* AA07_03 at 6 and 12 dpi; a 5-fold increase in bacterial biomass was observed for both Psa3 X-27 and Psa3 V-13 Δ*sEEL* relative to Psa3 V-13 using the qPCR approach (Figure 3A). Qualitatively, this same trend was also observed using the plate count method to quantify Psa biomass (Figure 3B). A linear correlation was observed when the dependent variables from the plate count (Log_10_ cfu/cm^2^) and qPCR (ΔCt) methodologies were plotted against one another as a regression analysis, specifically at 12 dpi (Figure 3C).

**Figure 3:**
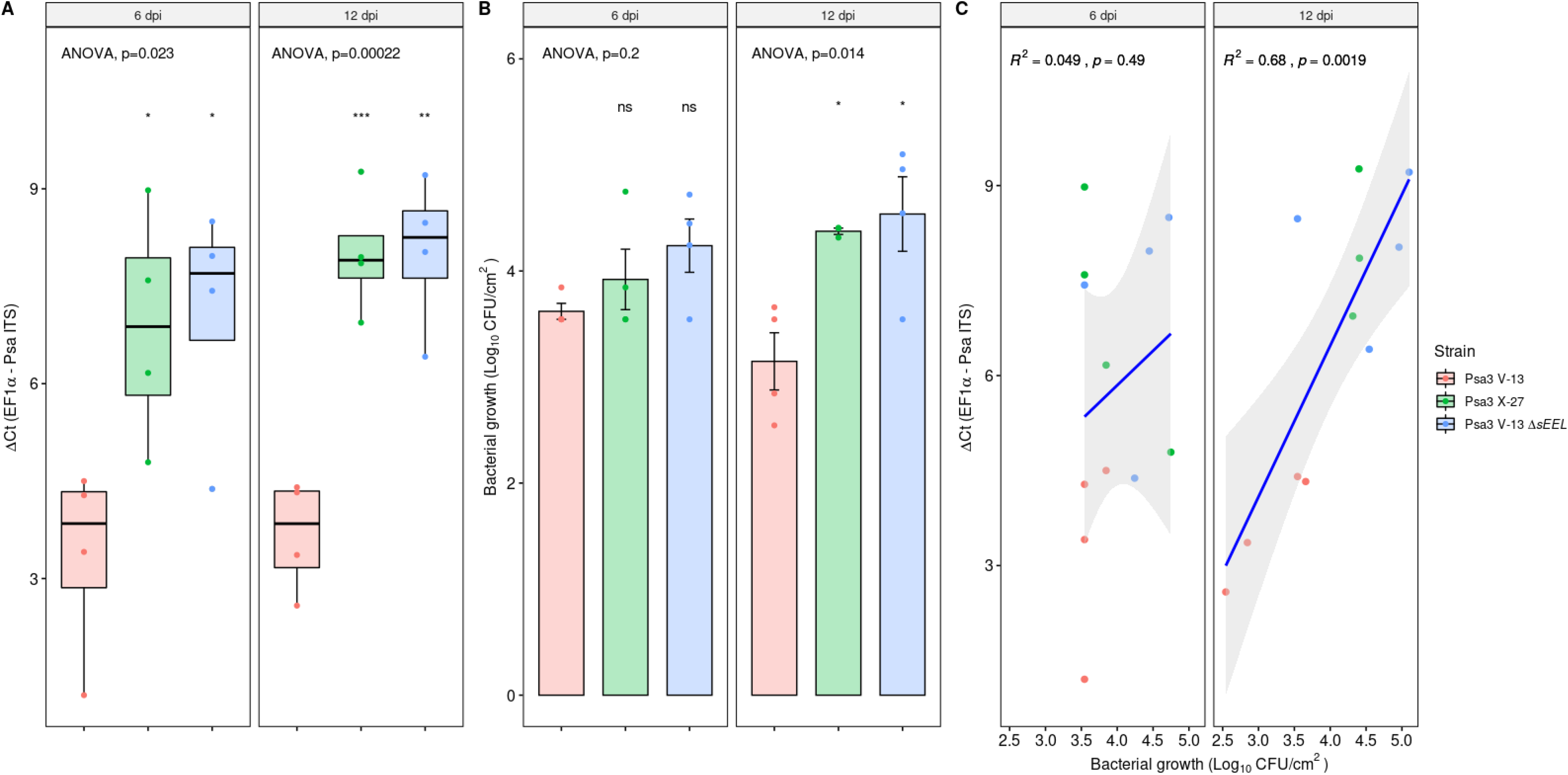
Psa3 X-27 and Psa3 V-13 Δ*sEEL* escape host recognition in *Actinidia arguta*. *A. arguta* AA07_03 plantlets were flood-inoculated with Psa3 V-13, Psa3 X-27, and Psa3 V-13 Δ*sEEL* at approximately 10^6^ cfu/mL. Bacterial growth was quantified at 6 and 12 days post-inoculation by qPCR ΔCt analysis **A** and plate count quantification **B**. **(A)** Box and whisker plots, with black bars representing the median values for the four pseudobiological replicates and whiskers representing the 1.5 inter-quartile range. **(B)** Bar height represents the mean number of Log_10_ cfu/cm^2,^ and error bars represent the standard error of the mean (SEM) between four pseudobiological replicates. **(C)** Regression analysis comparing the two quantification methods (**A** and **B**). The linear regression line is indicated in blue, the grey region indicates a 95% confidence interval, and the r-value represents the correlation coefficient (R^2^) and its associated p-value. The experiments were repeated three times with similar results. Asterisks indicate the statistically significant difference of Student’s *t*-test between the indicated strain and wild-type Psa3 V-13, where p≤.05 (*), p≤.01 (**), p≤.001 (***), and p>.05 (ns; not significant).

At 50 dpi, AA07_03 plantlets inoculated with Psa3 V-13 appeared healthy, with little to no development of disease symptoms (Figure S1). Conversely, AA07_03 plantlets inoculated with Psa3 X-27 developed leaf yellowing with small, angular, necrotic lesions surrounded by chlorotic halos. Similar disease-like symptoms were observed when AA07_03 was inoculated with Psa3 V-13 Δ*sEEL* (Figure S1). Quantification of diseased tissue (chlorotic and necrotic tissues) using a PIDIQ pipeline (Laflamme et al., 2016) indicated a clear difference between Psa3 V-13-infected versus Psa3 X-27- and Psa3 Δ*sEEL*-infected plants (Figure S2). Unlike AA07_03, *A. chinensis* var. *chinensis* ‘Hort16A’ is highly susceptible to Psa3 V-13. At 50 dpi, ‘Hort16A’ plantlets inoculated with Psa3 V-13 had a high degree of leaf yellowing and large areas of necrosis (Figure S1). Psa3 X-27 and Psa3 V-13 Δ*sEEL* both produced similar disease symptoms to Psa3 V-13 in ‘Hort16A’, with widespread necrosis evident (Figure S1).

### Four candidate avirulence effector loci contribute to Psa3 recognition in *A. arguta*

Knocking out the sEEL locus increased virulence in *A. arguta* AA07_03 quantitatively, but Psa3 X-27 or Psa3 V-13 Δ*sEEL* were not as virulent in *A. arguta* AA07_03 as they were in ‘Hort16A’. This suggested that there may be additional effectors recognized by AA07_03 within the Psa3 V-13 effector complement.

To determine whether additional Psa3 V-13 effectors triggered resistance in *A. arguta*, a library of 21 knockout strains was generated, covering all 30 effectors from Psa3 V-13, consisting of 15 individual effectors, a redundant effector pair (*hopAM1a-1*/*hopAM1a-2*), effector blocks (*hopZ5a*/*hopH1a*, CEL, or three different iterations of the EEL – Figure S3). This library of knockout strains was screened in *A. arguta* AA07_03 plantlets by flood-inoculation and sampled at 12 dpi (Figure 4). qPCR bacterial biomass quantification alone was used for this screen, owing to the large number of strains being assessed for pathogenicity across three independent infection experiments.

**Figure 4:**
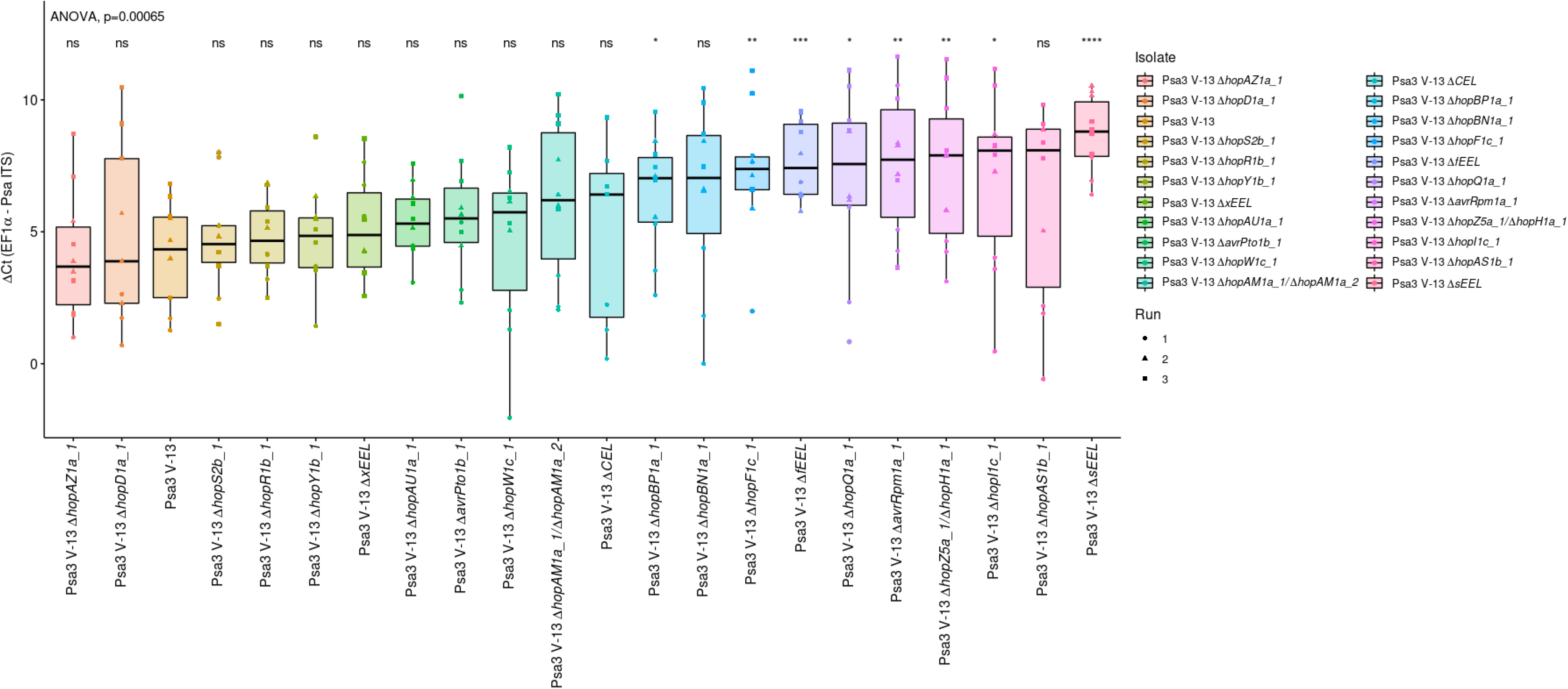
Pathogenicity assay screen of Psa3 V-13 effector knockout strains in *Actinidia arguta* identifies four avirulence loci. *A. arguta* AA07_03 kiwifruit plantlets were flood-inoculated at approximately 10^6^ cfu/mL. Psa biomass (*ITS*) was quantified relative to *AaEF1α* using the ΔCt analysis method for three pseudobiological replicates, per strain, per experimental run. Box and whisker plots, with black bars representing the median values and whiskers representing the 1.5 inter-quartile range. Asterisks indicate the statistically significant difference of Student’s *t*-test following ANOVA between the indicated strain and wild-type Psa3 V-13, where p ≤.05 (*), p≤.01 (**), p≤.001 (***), p≤.0001 (****), and p>.05 (ns; not significant). This experiment was separately conducted three times (biological replicates) with three batches of independently grown plants and data were stacked to generate the box plots.

Several effector knockout strains achieved significantly more bacterial growth than Psa3 V-13, including Psa3 V-13 Δ*sEEL*, partially escaping recognition in AA07_03 (Figure 4). Additionally, Psa3 V-13 Δ*fEEL,* which encompasses the sEEL effectors alongside additional effectors in the EEL (*avrB2b*, *avrD1*, and *hopF4a*), was also significantly more virulent in AA07_03 than Psa3 V-13 (Figure 4). Conversely, Psa3 V-13 Δ*xEEL*, which encompasses the fEEL effectors alongside additional effectors in the EEL (*hopQ1a* and *hopD1a*), and Psa3 V-13 Δ*CEL* were not significantly different from Psa3 V-13. Psa3 V-13 Δ*hopZ5a/*Δ*hopH1a*, Psa3 V-13 Δ*avrRpm1a* and Psa3 V-13 Δ*hopF1c* also had a significant increase (p ≤ 0.01) in bacterial growth *in planta* relative to Psa3 V-13. The isolates Δ*hopI1c*, Δ*hopBP1a*, and Δ*hopQ1a* had a significant increase (p ≤ 0.05) and Δ*hopBN1a* was not significant overall but was significant in two of the three qPCR runs. These were further tested by plate count methods and were not found to be significantly increased in virulence in AA07_03 compared with Psa3 V-13 (Figure S4).

Following this screen, candidate avirulence effector knockout strains with significance (p<0.01) were tested by plate count methods. Using the previously described biolistic co-expression assays to measure HR-mediated reporter eclipse in AA07_03 leaves, *hopZ5a* was identified as the recognized effector in the *hopZ5a*/*hopH1a* effector block (Figure S5; (Jayaraman et al., 2021). Therefore, only the single *hopZ5a* knockout strain was used for subsequent experiments. Thus, the candidate avirulence-effector knockout strains selected for further analysis were Psa3 V-13 Δ*sEEL*, Psa3 V-13 Δ*fEEL*, Psa3 V-13 Δ*xEEL*, Psa3 V-13 Δ*hopZ5a*, Psa3 V-13 Δ*hopF1c*, and Psa3 V-13 Δ*avrRpm1a*. Psa3 V-13 Δ*hopI1c* was selected to be a negative control in this experiment, as this strain did not display an increase in bacterial growth or escape recognition because of the deletion of the Δ*hopI1* effector gene (Figure S4). To confirm the candidate avirulence effector knockout strains identified in the qPCR screens, bacterial growth was quantified in AA07_03 using both qPCR and the plate count method (Figures 5A and 5B; Figures S6 and S7). Interestingly, all three of the *EEL* knockout strains had significantly more Psa biomass *in planta,* with a ten-fold increase in bacterial growth relative to Psa3 V-13. Similarly, Psa3 V-13 Δ*hopZ5a,* Psa3 V-13 Δ*hopF1c* and Psa3 V-13 Δ*avrRpm1a* also had significantly more bacterial growth *in planta* relative to Psa3 V-13, with approximately a mean ten-fold increase in bacterial growth. As expected, Psa3 V-13 Δ*hopI1c* was not significantly different from Psa3 V-13.

**Figure 5:**
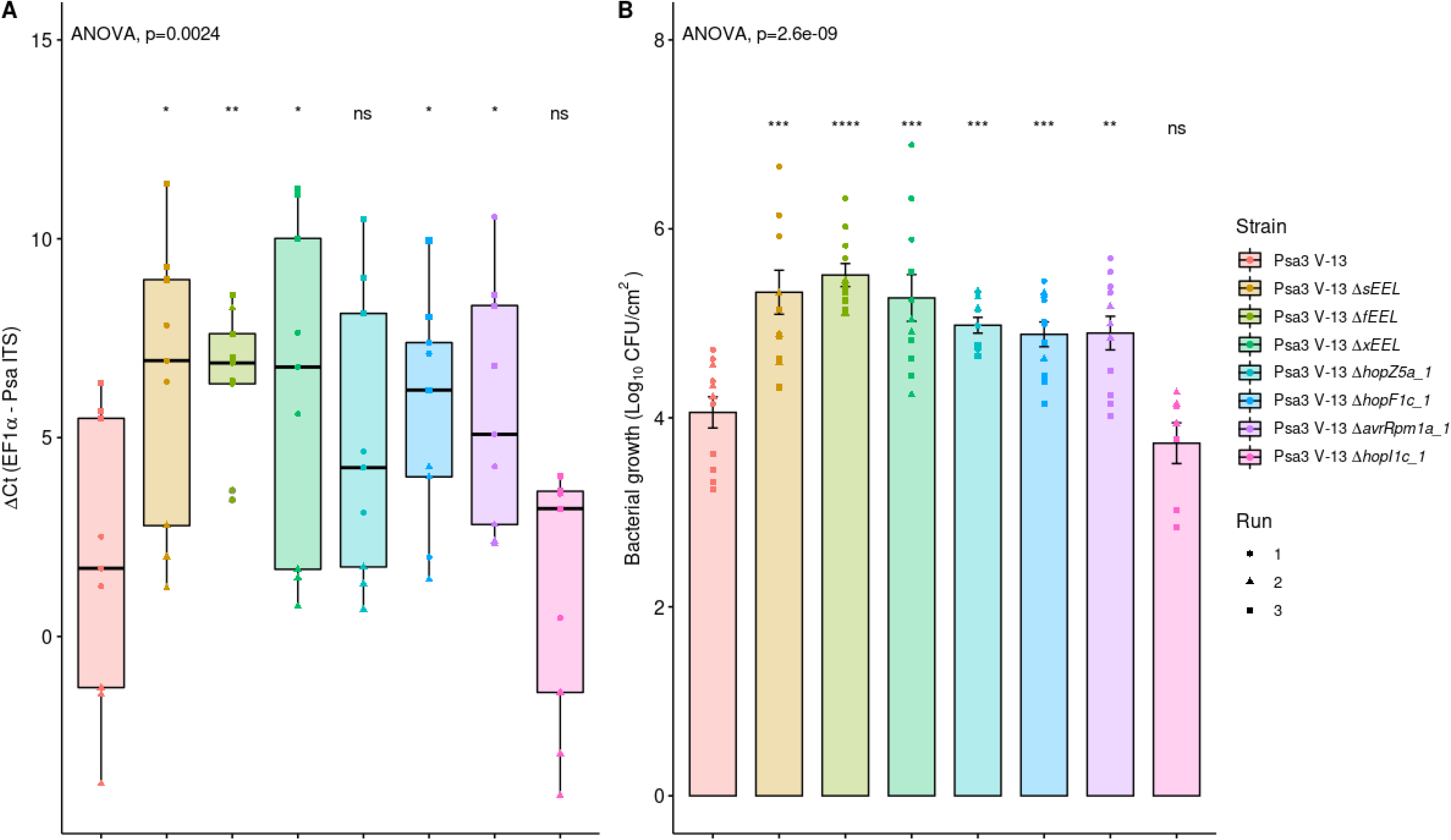
Pathogenicity assay of Psa3 V-13 selected effector knockout strains in *Actinidia arguta* confirming four loci recognition. *A. arguta* AA07_03 kiwifruit plantlets were flood-inoculated at approximately 10^6^ cfu/mL. Bacterial pathogenicity was quantified relative to Psa3 V-13 using the ΔCt analysis method and box and whisker plots, with black bars representing the median values and whiskers representing the 1.5 inter-quartile range **(A)** and plate count quantification and error bars representing the standard error of the mean (SEM) **(B),** for four pseudobiological replicates, per strain, per experimental run. Asterisks indicate the statistically significant difference of Student’s *t*-test between the indicated strain and wild-type Psa3 V-13, where p ≤.05 (*), p≤.01 (**), p≤.001 (***), and p>.05 (ns; not significant). This experiment was separately conducted three times (biological replicates) with three batches of independently grown plants and data were stacked to generate the box plots and bar graphs shown.

### sEEL effector HopAW1a triggers resistance in *A. arguta*

Pathogenicity screening of AA07_03 determined that Psa3 V-13 Δ*sEEL* lost at least one avirulence effector (Figure 5). To identify which sEEL effector(s) triggers resistance, individual sEEL effectors (*hopAW1a*, *hopD2a*, *hopF1e* and *hopAF1b*) were plasmid-complemented into Psa3 V-13 Δ*sEEL* (Table 1). Pathogenicity assays were conducted to identify which sEEL effector(s) triggered resistance in *A. arguta* AA07_03.

**Table 1.**
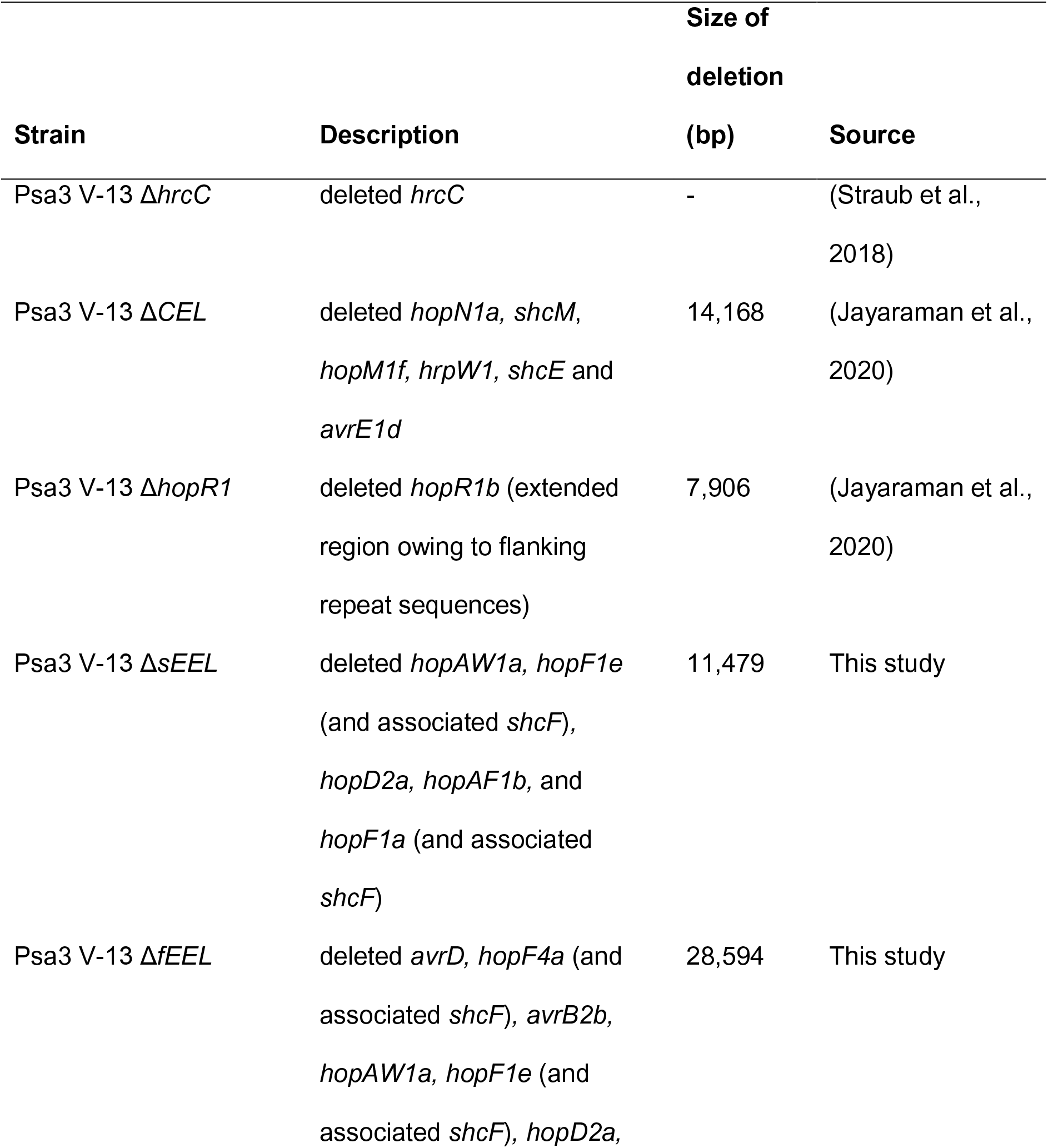
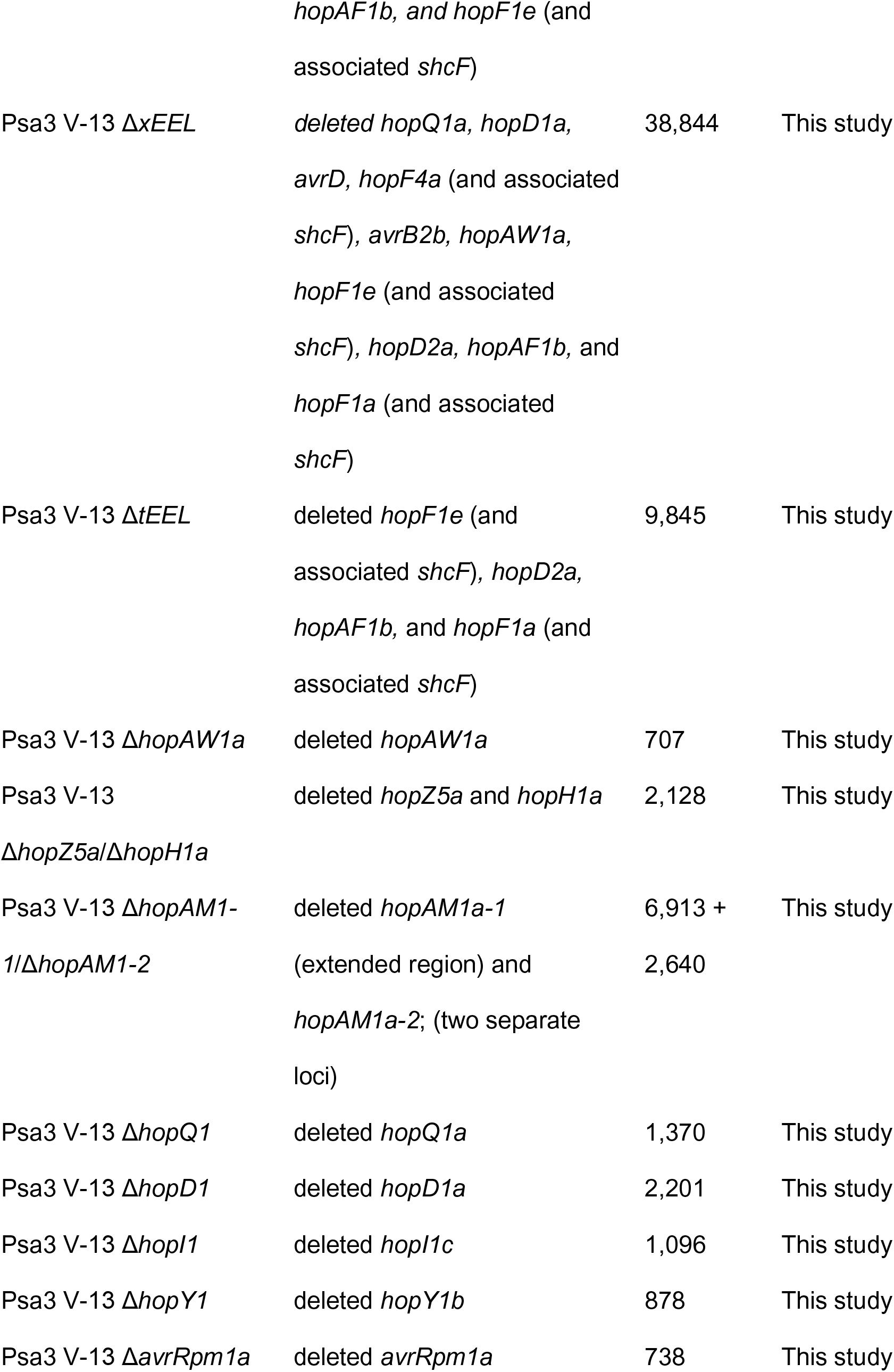
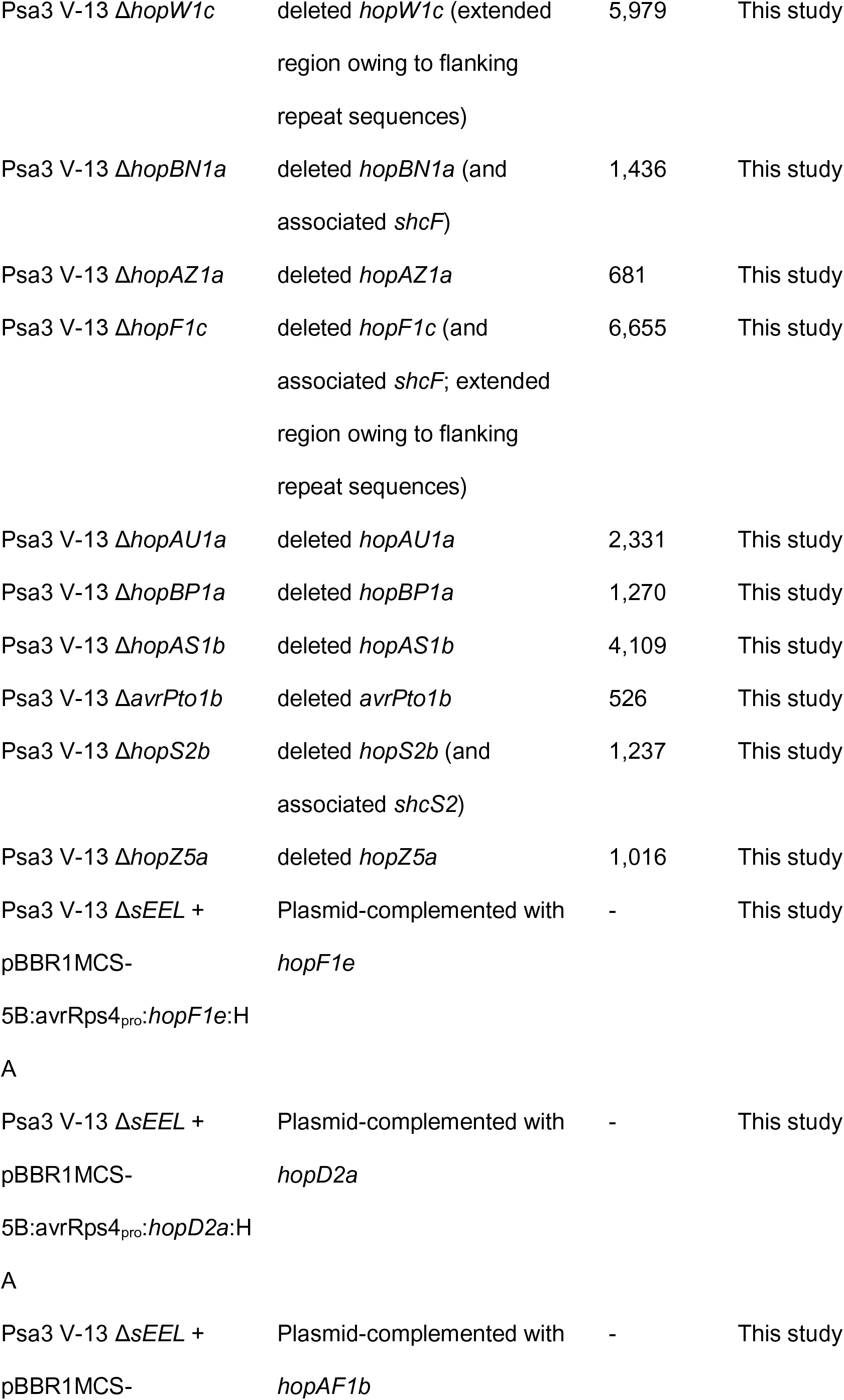
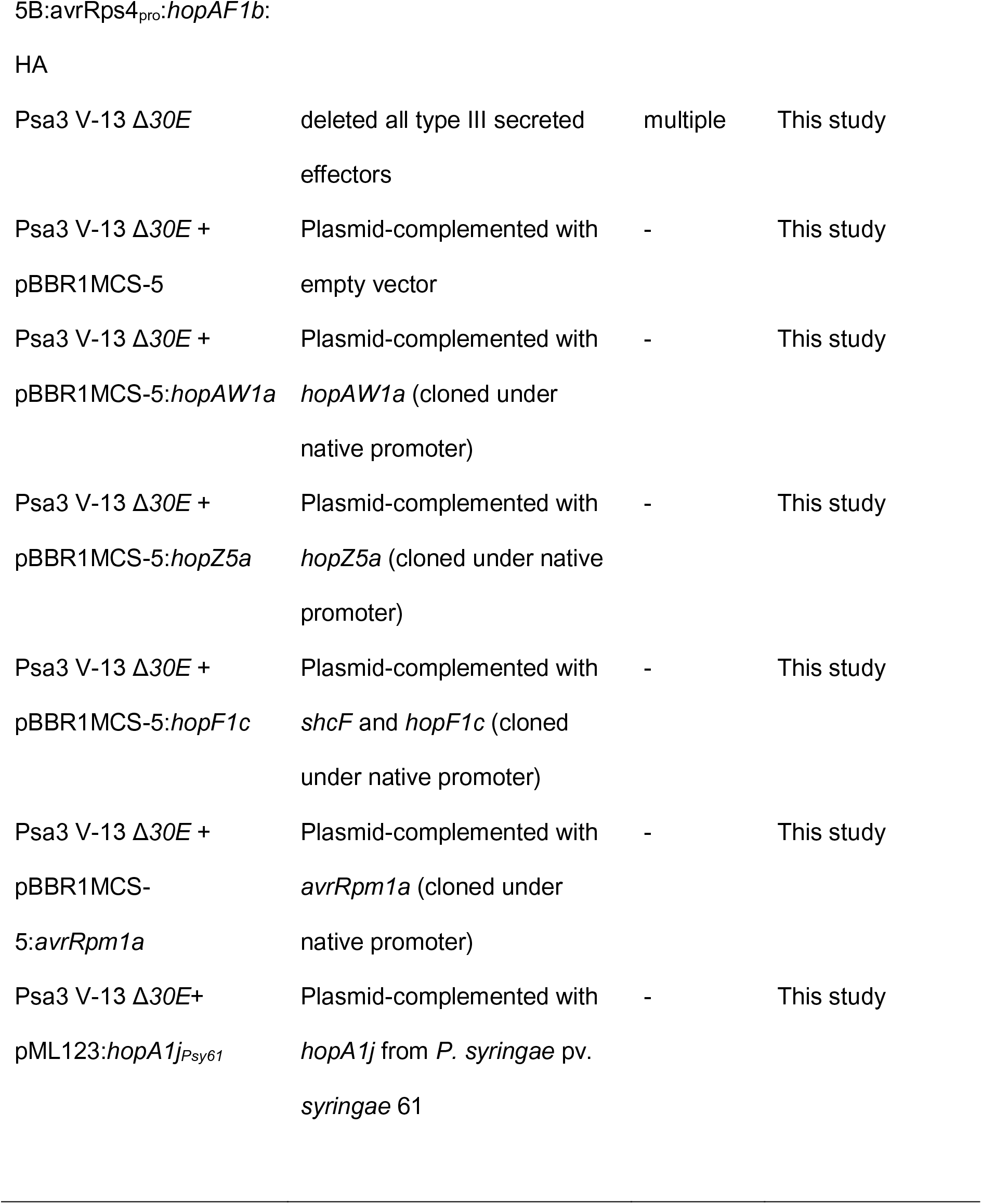
Transgenic Psa3 V-13 effector knockout and plasmid-complemented strains.

Plasmid complementation of Psa3 V-13 Δ*sEEL* with *hopAF1b* and *hopD2a* yielded similar amounts of *in planta* bacterial biomass to Psa3 V-13 Δ*sEEL* and these were significantly different from Psa3 V-13 (Figures 6A and 6B). This suggests that neither HopAF1b nor HopD2a trigger resistance to Psa3 V-13 in AA07_03. Interestingly, Psa3 V-13 Δ*sEEL* + *p.hopAW1a* and Psa3 V-13 Δ*sEEL* + *p.hopF1e* showed a decrease in *in planta* bacterial biomass relative to Psa3 V-13 Δ*sEEL,* suggesting that individual plasmid complementation of *hopAW1a* and *hopF1e* partially restored host recognition (Figures 6A and 6B). However, using the qPCR method (Figure 6A), *in planta* bacterial biomass of neither of these strains was fully reduced to the same degree as Psa3 V-13, possibly owing to plasmid loss. If both effectors are required for recognition, they may have an additive effect that is only fully seen in wild-type Psa3 V-13. The plate count quantification (Figure 6B), in contrast, showed neither Psa3 V-13 Δ*sEEL* + *p.hopAW1a* nor Psa3 V-13 Δ*sEEL* + *p.hopF1e* was significantly different from Psa3 V-13, suggesting that both HopAW1a and HopF1e may trigger resistance in AA07_03.

**Figure 6:**
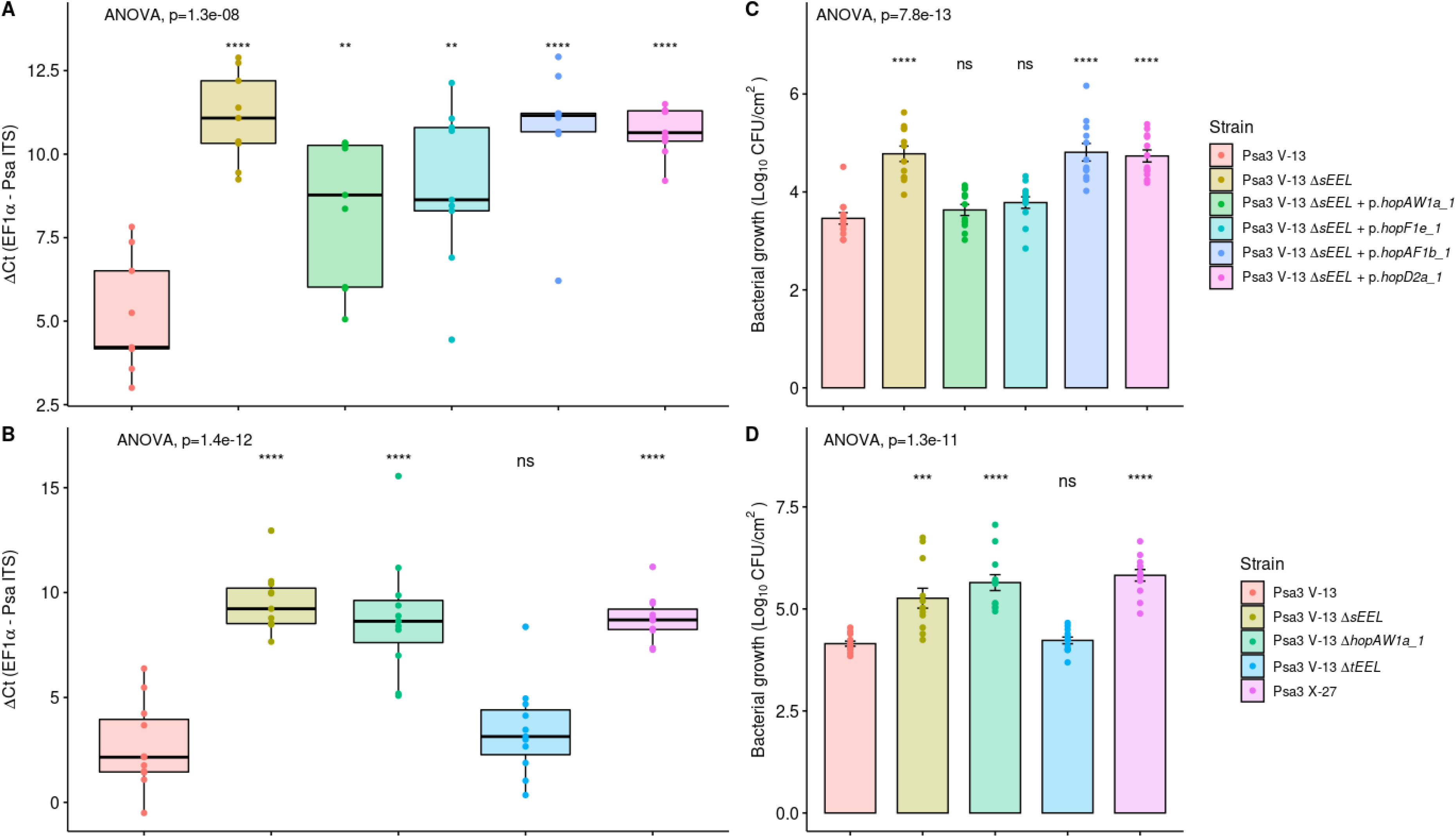
Psa3 V-13 *EEL* effector HopAW1a is recognized by *Actinidia arguta*. *A. arguta* AA07_03 plantlets were flood-inoculated at approximately 10^6^ cfu/mL. Bacterial growth was quantified 12 days post-inoculation using qPCR ΔCt analysis **A,** and plate count quantification **B,** for the plasmid-complemented *ΔsEEL* strains. Asterisks indicate the statistically significant difference of Student’s *t*-test between the indicated strain and wild-type Psa3 V-13, where p≤0.01 (**), p≤0.0001 (****), p>.05 (ns). **(A)** Box and whisker plots, with black bars representing the median values and whiskers representing the 1.5 inter-quartile range. **(B)** Bar height represents the mean number of Log_10_ cfu/cm^2,^ and error bars represents the standard error of the mean (SEM) between four pseudobiological replicates. Bacterial growth was quantified 12 days post-inoculation using qPCR ΔCt analysis **C,** and plate count quantification **D,** for the *ΔtEEL* and *ΔhopAW1a* strains. Asterisks indicate the statistically significant difference of Student’s *t*-test between the indicated strain and wild-type Psa3 V-13, where p≤.001 (***), p≤.0001 (****), and p>.05 (ns). **(C)** Box and whisker plots, with black bars representing the median values and whiskers representing 1.5 inter-quartile range. **(D)** Bar height represents the mean number of Log_10_ cfu/cm^2,^ and error bars represents the standard error of the mean (SEM) between four pseudobiological replicates. Both experiments (**A**, **B** and **C**, **D**) were separately conducted three times (biological replicates) with three batches of independently grown plants and data were stacked to generate the box plots and bar graphs shown.

To confirm that HopAW1a and HopF1e are candidate avirulence effectors, segmented effector knockouts within the sEEL were generated to confirm these results (Figures 6C and 6D). Psa3 V-13 Δ*hopAW1a* lacks *hopAW1a* while Psa3 V-13 Δ*tEEL* lacks *hopF1e*, *hopAF1b*, *hopD2a* and *hopF1a*. Pathogenicity assays of AA07_03 demonstrated that Psa3 V-13 Δ*hopAW1a* was significantly different from Psa3 V-13 and similar to Psa3 V-13 Δ*sEEL*. In contrast, Psa3 V-13 Δ*tEEL* was not significantly different from Psa3 V-13. This suggests that the individual deletion of *hopAW1a* is sufficient to partially release host recognition and further suggests that none of the effectors in the tEEL triggers resistance on AA07_03. The plate count data (Figure 6D) results corroborate the qPCR data (Figure 6C) and suggest that HopAW1a is the sole sEEL effector responsible for triggering resistance on AA07_03. Notably, AA07_03 plantlets inoculated with Psa3 Δ*sEEL* complemented with *hopAW1a* was the sole plasmid-complemented line to display a lack of disease symptoms, including leaf yellowing and necrosis (Figure S8). Additionally, Psa3 Δ*hopAW1a* produced Psa3 Δ*sEEL*-like disease symptoms while Psa3 Δt*EEL* did not (Figure S8). Quantification of diseased tissue (chlorotic and necrotic tissues) using the modified PIDIQ pipeline indicated Psa3 Δ*sEEL*-infected plants most closely resembled the Psa3 Δ*hopAW1a-*infected plants, while Psa3 V-13-infected plants resembled Psa3 Δ*tEEL*-infected plants (Figure S9). These results were further supported by biolistic co-expression assays in AA07_03 leaves, with only *hopAW1a* triggering an HR response and an associated reporter eclipse (see below, Figure S10).

### Psa3 candidate avirulence effectors trigger a hypersensitive response in *A. arguta*

Psa3 V-13 effectors *hopAW1a, hopF1c, hopZ5a,* and *avrRpm1a* cloned under a 35S promoter were co-bombarded into kiwifruit leaf tissue with a GUS reporter gene to assess if the proteins they encode triggered the hypersensitive response (HR) in *A. arguta* AA07_03 and *A. chinensis* var. *chinensis* ‘Hort16A’ leaves (Figure 7A). The effector *hopA1j* from *P. syringae* pv. *syringae* 61 was used as a positive control for HR in this assay (Jayaraman et al., 2021). Co-bombardment of candidate avirulence effectors *hopAW1a, hopF1c, hopZ5a* and *avrRpm1a* all demonstrated a decrease in GUS activity on *A. arguta* AA07_03 in comparison to the control (empty vector), indicating that the proteins they encode triggered a hypersensitive response. Surprisingly, HopF1c expression in ‘Hort16A’ leaves also produced an HR similar to that in AA07_03, albeit without a significant difference in ion leakage compared with the control. The HR triggered by AvrRpm1a, HopZ5a and HopAW1a appeared to be AA07_03-specific, however. Ion leakage assays using *Pseudomonas fluorescens* Pf0-1 carrying an introduced type III secretion system indicated that only HopAW1a resulted in an increase in conductivity compared with the empty vector control, although the result was considerably weaker than that with the control HopA1j (Figure S11).

**Figure 7:**
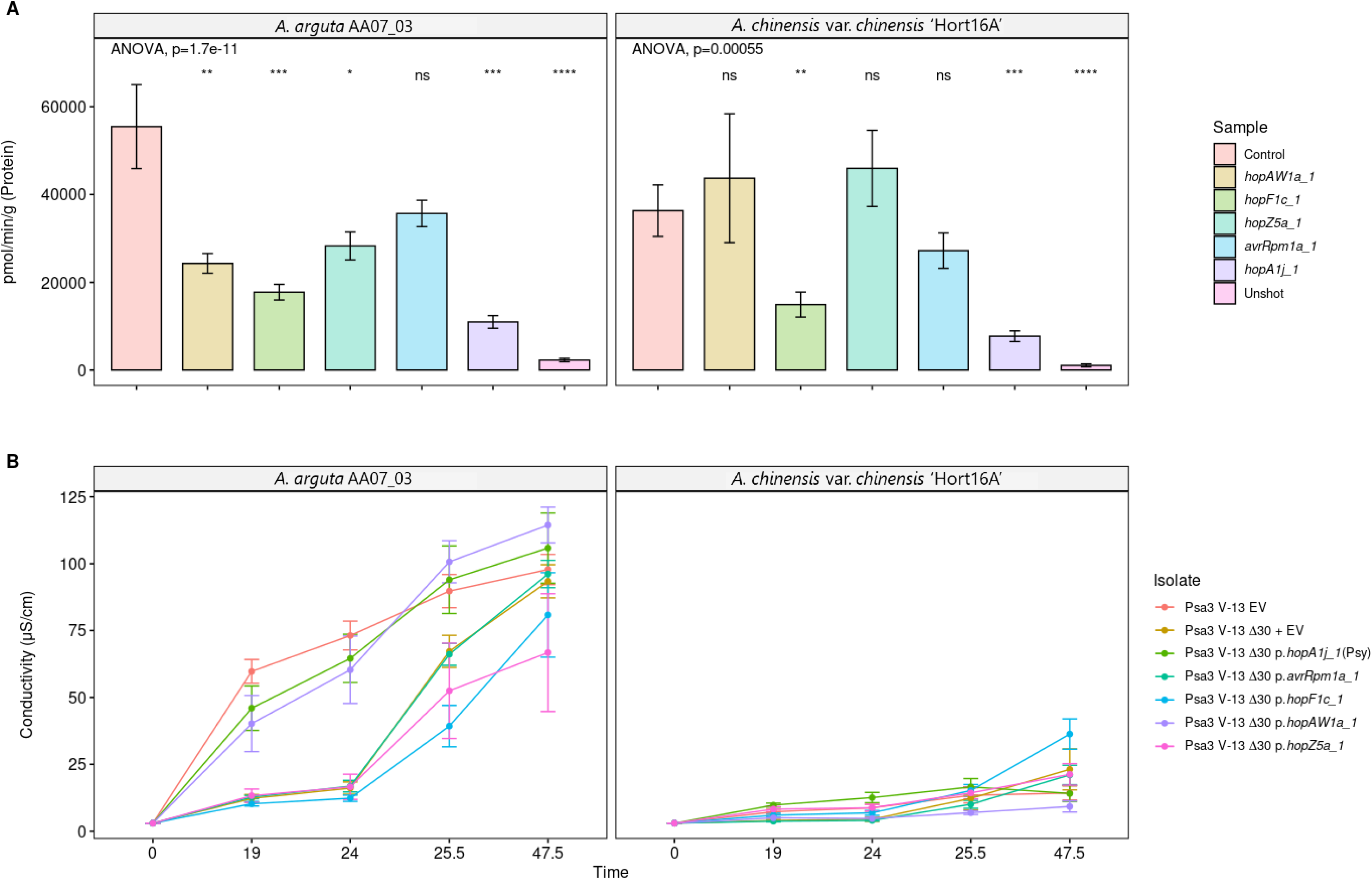
Reporter eclipse assays demonstrate that HopAW1a, HopZ5a, and AvrRpm1a trigger a host-specific immunity response in *Actinidia arguta* partially supported by ion leakage assays. **(A)** Avirulence effectors cloned in binary vector constructs tagged with GFP, or an empty vector (Control), were co-expressed with a β-glucuronidase (GUS) reporter construct using biolistic bombardment and priming in leaves from *A. arguta* AA07_03 or *A. chinensis* var. *chinensis* ‘Hort16A’ plantlets (Jayaraman et al., 2021). The GUS activity was measured 48 hours after DNA bombardment. Error bars represent the standard errors of the means for three independent biological replicates with six technical replicates each (n=18). HopA1j from *Pseudomonas syringae* pv. *syringae* 61 was used as positive control and un-infiltrated leaf tissue (Unshot) as a negative control. Tukey’s HSD indicates treatment groups which are significantly different at α ≤ 0.05 with different letters. **(B)** Leaf discs from *A. arguta* AA07_03 and *A. chinensis* var. *chinensis* ‘Hort16A’ plantlets were vacuum-infiltrated with Psa3 V-13 wild-type strain or Psa3 V-13 Δ*30E* carrying empty vector (EV), or Psa3 V-13 Δ*30E* carrying a plasmid-borne type III secreted effector (*hopAW1a*, *hopZ5a*, *avrRpm1a* or *shcF*:*hopF1c*, or positive control *hopA1j* from *P. syringae* pv. *syringae* 61) inoculum at ∼5 x 10^8^ cfu/mL. Electrical conductivity due to HR-associated ion leakage was measured at indicated times over 72 hours. The ion leakage curves are faceted by plant species and stacked for three independent runs of this experiment. Error bars represent the standard errors of the means calculated from the five pseudobiological replicates per experiment (n=15).

*P. fluorescens* may not be able to express and deliver effectors from Psa in the full context of a suite of other potential pathogenicity factors. To mimic this more complete context and deliver individual effectors from Psa3, a complete effector knockout strain (Psa3 V-13 Δ*30E*) was generated that lacked all 30 predicted and expressed effectors. To confirm that the AA07_03-recognized avirulence effectors were not affected by level of expression under the synthetic promoter or the C-terminal HA tag, *hopAW1a*, *hopZ5a*, *avrRpm1a*, and *hopF1c* (with *shcF* carrying a point mutation resulting in an early truncation) were cloned under their native promoters. Psa3 V-13 Δ*30E* delivery of *hopAW1a*, *hopZ5a*, *avrRpm1a*, and *hopF1c* surprisingly revealed that only HopAW1a was able to trigger a strong early ion leakage in AA07_03 leaves (24 h), but not in ‘Hort16A’ (Figure 7B). Owing to the lack of a functional ShcF protein, unsurprisingly, HopF1c only triggered a delayed ion leakage, but did so in both plants, supporting the reporter eclipse findings (Figure 7A).

### Cumulative deletion of Psa3 candidate avirulence effectors does not result in added fitness in *A. arguta*

To identify whether Psa3 V-13 avirulence effectors *hopF1c*, *avrRpm1a*, *hopZ5a*, and *hopAW1a* contribute cumulatively towards triggering resistance, all four effectors were successively knocked out of the Psa3 V-13 strain and these multiple-knockout strains were inoculated onto *A. arguta* AA07_03 and *A. chinensis* var. *chinensis* ‘Hort16A’ plantlets (Figure 8). Psa3 V-13 Δ*hrcC* was used as a negative control, as it lacks the ability to secrete type III effectors into host plant cells and is not virulent in *Actinidia* host plants, including ‘Hort16A’.

**Figure 8:**
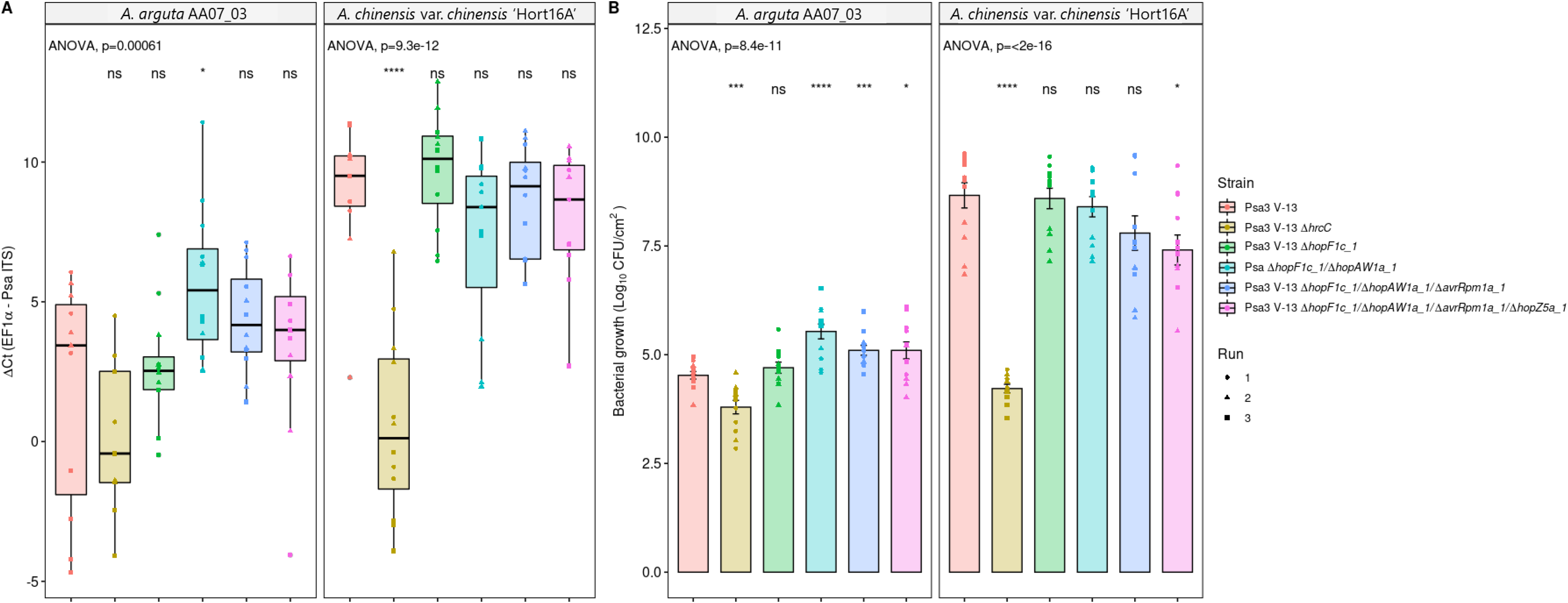
Pathogenicity assay of Psa3 V-13 multiple avirulence effector knockout strains demonstrates lack of increasing resistance-escape due to a cumulative loss of virulence. *Actinidia arguta* AA07_03 and *Actinidia chinensis* var. *chinensis* ‘Hort16A’ kiwifruit plantlets were flood-inoculated at approximately 10^6^ cfu/mL. Bacterial growth was quantified at 12 days post-inoculation using qPCR ΔCt analysis **A** and plate count quantification **B**. The experiment was conducted three times (biological replicates) with three batches of independently grown plants and data were stacked to generate the box plots and bar graphs shown. Asterisks indicate significant differences from ANOVA followed by a *post hoc* Student’s *t*-test between the indicated strain and wild-type Psa3 V-13, where p ≤005 (*), p≤.001 (***), p≤.0001 (****), and p>.05 (ns; not significant). **(A)** Box and whisker plots, with black bars representing the median values, whiskers representing the 1.5 inter-quartile range, and black dots indicating outliers. **(B)** Bar height represents the mean number of Log_10_ cfu/cm^2^ and error bars represents the standard error of the mean (SEM) between four pseudobiological replicates.

Interestingly, while Psa3 V-13 is avirulent in AA07_03, the type III secretion-deficient mutant (Psa3 V-13 Δ*hrcC*) grew less than the wild-type, suggesting that while several effectors trigger a strong HR in AA07_03 plants, the retention of effector secretion remains largely beneficial to Psa3 (Figures 8A and 8B). Furthermore, pathogenicity assays in AA07_03 demonstrated that, while Psa3 V-13 Δ*hopF1c*/Δ*hopAW1a* (double), Psa3 V-13 Δ*hopF1c*/Δ*hopAW1a*/Δ*avrRpm1a* (triple), and Psa3 V-13 Δ*hopF1c*/Δ*hopAW1a*/Δ*avrRpm1a*/Δ*hopZ5a* (quadruple) were significantly different from Psa3 V-13, they did not cumulatively increase in growth *in planta* with each successive knockout (Figures 8A and 8B). In fact, the triple and quadruple effector knockout strains appeared to accumulate in reduced amounts in AA07_03 compared with the double knockout. This finding of reduced fitness in AA07_03 for the multiple knockout strains was largely reflected in ‘Hort16A’, with the quadruple knockout demonstrating nearly 15-fold less growth compared with Psa3 V-13 (Figure 8A). Taken together, the data suggest that while several Psa3 effectors are recognized in *A. arguta*, the ability to secrete these effectors collectively is beneficial to survival in kiwifruit plants and thus they are unlikely to be lost in succession from a lack of evolutionary selection.

### Psa3 avirulence effectors shared by multiple Psa biovars appear to contribute to broad Psa resistance in *A. arguta*

The four Psa3 V-13 effectors we have identified that are recognized in *A. arguta* AA07_03 are also present in the effector complements of the other Psa biovars. At least one avirulence effector is shared for each emergent clade of Psa with *hopAW1a* in Psa5/Psa6, *avrRpm1a* in Psa1/Psa6, and *hopF1c* in Psa2/Psa5 (Figure 9A). Because Psa2 possesses a close orthologue of a truncated effector in Psa3 V-13 (*avrRpm1c*), we checked whether AvrRpm1c was also recognized in AA07_03 and ‘Hort16A’ leaves. Similar to AvrRpm1a, AvrRpm1c from Psa2 K-28 was also recognized specifically in AA07_03 but not in ‘Hort16A’ (Figure S12).

**Figure 9:**
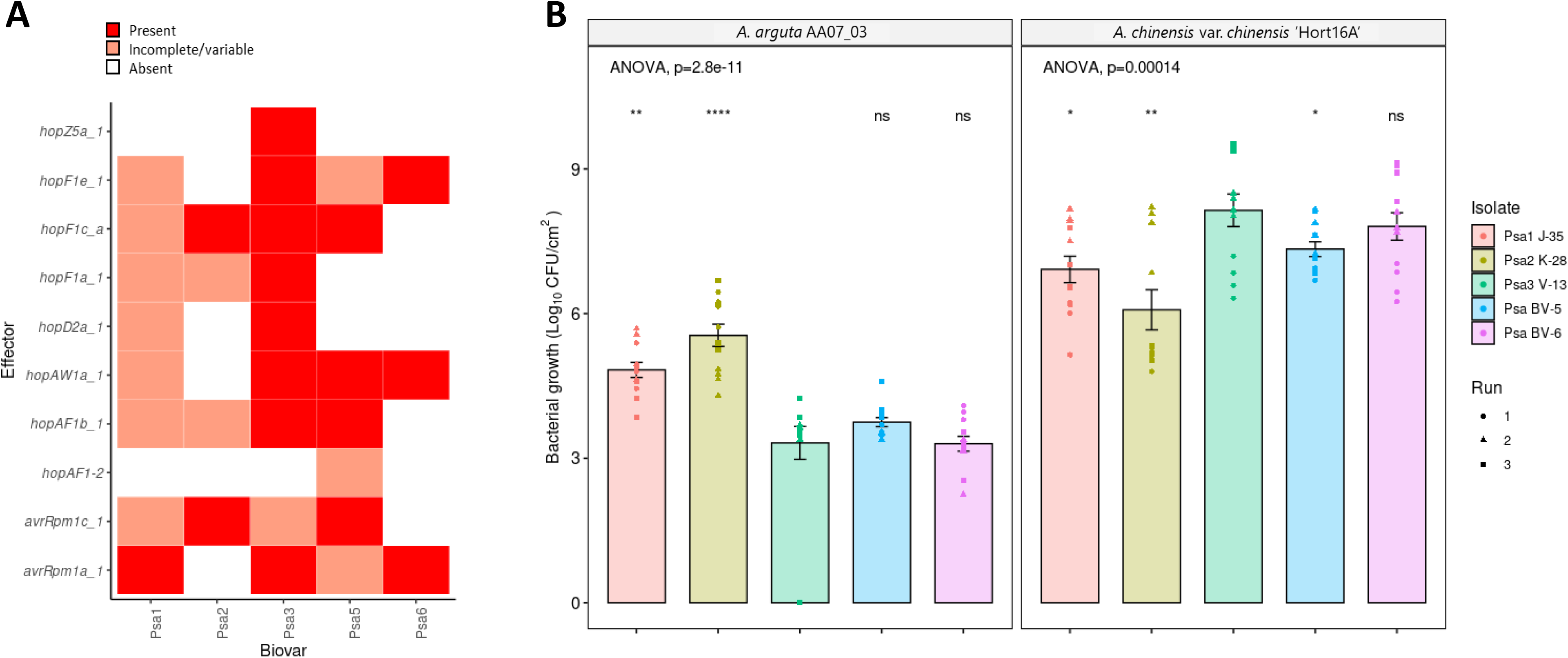
Pathogenicity assay of *Pseudomonas syringae* pv. *actinidiae* (Psa) biovars in *Actinidia arguta* indicates broad recognition across biovars. **(A)** Effectors of interest across the Psa biovars. Selected effector repertoires collated from McCann et al. (2013) and Sawada et al. (2016). Red indicates when an effector is present; pink indicates when an effector is either incomplete or variable, i.e. present within some isolates of the biovar but not others; and white indicates when an effector is absent from a given biovar. **(B)** *A. arguta* AA07_03 and *A. chinensis* var. *chinensis* ‘Hort16A’ kiwifruit plantlets were flood-inoculated at approximately 10^6^ cfu/mL with Psa1 J-35, Psa2 K-28, Psa3 V-13, Psa5 MAFF212057, and Psa6 MAFF212134 strains. Bacterial growth was quantified at 12 days post-inoculation using plate count quantification. The experiment was conducted three times (biological replicates) with three batches of independently grown plants and data were stacked to generate the box plots and bar graphs shown. Asterisks indicate significant differences from ANOVA followed by a *post hoc* Student’s *t*-test between the indicated strain and wild-type Psa3 V-13, where p≤.05 (*), p≤.01 (**), p≤.001 (***), p≤.0001 (****), and p>.05 (ns; not significant). Bar height represents the mean number of Log_10_ cfu/cm^2^ and error bars represents the standard error of the mean (SEM) between four pseudobiological replicates.

Having examined the effector complement of Psa3 V-13, we next sought to examine whether the presence of these shared avirulence effectors predicted performance of these biovars in *A. arguta* AA07_03. Representative Psa biovar strains were screened in *A. chinensis* var. *chinensis ‘*Hort16A’ and *A. arguta* AA07_03 to test their virulence (Figure 9B). At 12 dpi, the bacterial growth of Psa1 J-35, Psa2 K-28, and Psa5 in *A. chinensis* var. *chinensis* ‘Hort16A’ was slightly but significantly lower than that of Psa3 V-13, while that of Psa6 was not significantly different (Figure 9B). Conversely, in *A. arguta* AA07_03, Psa1 J-35 and Psa2 K-28 accumulated in significantly higher amounts than Psa3 V-13 at 12 dpi. Similarly, Psa5 accumulated in slightly higher amounts than Psa3 V-13 at 12 dpi, albeit not significantly. Meanwhile, Psa6 accumulated *in planta* in amounts similar to those of Psa3 V-13. This relationship between Psa growth in *A. arguta* and *A. chinensis* var. *chinensis* appeared to be inversely correlated. These results taken together suggest a broad recognition, present specifically in *A. arguta*, of a number of shared effectors across the Psa biovars.

## Discussion

Although the pandemic isolate Psa3 causes devastating leaf spot and canker symptoms in widely grown commercial cultivars of *A. chinensis*, many other *Actinidia* species such as *A. arguta* are resistant (Datson et al., 2013). Resistance to Psa3 in the *A. arguta* accessions we investigated was due to the induction of the hypersensitive response (HR). This conclusion was supported by both microscopic observation of cell death and ion leakage experiments using leaf discs (Jayaraman et al., 2021) (Figure 1). The induction of HR implies effector-triggered immunity (ETI), and that at least one effector in Psa3 is recognized by the product of an, as yet, uncharacterized, resistance gene in *A. arguta*. It is well documented that the loss of a single effector by deletion or mutation can increase virulence in previously resistant cultivars or species. Psa was isolated from leafspots of *A. arguta* plants in areas where Psa3 is prevalent in New Zealand. Whole genome sequence analysis of a Psa strain isolated from *A. arguta*, Psa3 X-27, identified a 49 kb deletion in the EEL, which included five effectors and the uncharacterized NRPS. This deletion appeared to be the only mutation of significance in these isolates. Flanking the deletion were two MITE sequences, suggesting a relatively facile mechanism for excision of the region via homologous recombination between the MITE loci.

A gene knock out in Psa3 V-13 that deleted the same group of five effectors present in Psa3 X-27 was constructed (Psa3 V-13 Δ*sEEL*). Both these isolates were able to grow to the same extent in AA07_03. This suggested that the deletion of the putative NRPS toxin biosynthesis gene cluster in Psa3 X-27 is not contributing to the increase in *in planta* growth. Several lines of evidence suggest that the increase in bacterial biomass associated with *A. arguta* infections by this strain is due only to the deletion of *hopAW1a*. These include: Psa3 V-13 *ΔsEEL* strain plasmid-complemented with *hopAW1a* demonstrated a decrease in pathogenicity to the same rate as that of Psa3 V-13; the Psa3 V-13 *ΔhopAW1a* individual effector knockout demonstrated an increase in pathogenicity similar to that observed with Psa3 X-27 and Psa3 V-13 *ΔsEEL*; and finally, biolistic expression of HopAW1a in AA07_03 leaves triggered an HR.

To determine whether there were additional effectors in Psa3 in addition to HopAW1a that trigger an HR, or whose loss might result in an increase in virulence in *A. arguta*, we generated and screened a library of effector knockouts for their ability to grow in AA07_03. We found that the loss of Psa3 V-13 effectors *avrRpm1a*, *hopF1c*, and *hopZ5a* increased growth in AA07_03 compared with Psa3 V-13, and biolistic expression of these effectors also indicated that they trigger an HR in *A. arguta*. Notably, this approach is unable to identify effectors that non-redundantly participate in virulence but are also recognized in *A. arguta*. Important Psa effectors that may fall into this category include AvrE1d and HopR1b (Jayaraman et al., 2020).

For the multiple candidate avirulence effectors identified, AA07_03 must either possess an R protein capable of recognizing numerous effectors, or carry multiple R proteins specific for each effector. If these avirulence effectors are collectively recognized by one or more host resistance proteins, the increase in bacterial growth observed in individual effector knockouts may not represent a full escape from host recognition. The complex interplay of effector complements makes it challenging to study the activity of a single effector in isolation. Modular co-expression of *Pseudomonas syringae* pv. *tomato* (Pto) DC3000 effectors has identified multiple instances of effector interplay; for example, the effector AvrPtoB is a suppressor of HopAD1-elicited ETI in *Nicotiana benthamiana* (Wei et al., 2018). Similarly, HopI1 can suppress ETI elicited by HopQ1-1 (Wei et al., 2018). Along these same lines, there may be further effectors that could be recognized by plant resistance proteins that have not been identified in this study owing to suppression of ETI by another effector. Suppression of ETI would prevent a decrease of bacterial biomass in its presence and, therefore, upon deletion we may not detect a change in bacterial biomass either.

Another complicating factor is redundancy – many effectors are collectively essential but individually redundant and can be grouped into redundant effector groups (REGs) (Kvitko et al., 2009). Redundancy can exist on numerous levels – redundant effectors may modify the same host target using the same molecular mechanism, or through different molecular mechanisms towards the same means of plant immunity suppression (Ghosh and O’Connor, 2017). Pto DC3000 possesses at least two REGs with redundant functions (Wei et al., 2018). One REG contains the *CEL* effectors *hopM1* and *avrE1*, which are redundantly involved in water-soaking in the apoplast, promoting bacterial growth (Wei et al., 2018). Another REG contains *avrPto* and *avrPtoB,* which redundantly suppress PTI induced by FLS2 perception of bacterial flagellin (Wei et al., 2018). Redundancy between effectors, alongside potential effector interplay, means that avirulence effector knockouts may not show significant changes in bacterial biomass if other avirulence effectors are still present and act epistatically.

One interesting commonality between the avirulence effectors identified in this research is that orthologs of several of these effectors from other *P. syringae* pathovars, including AvrRpm1 and HopF2, target the RIN4 plant defence signaling hub (Ray et al., 2019). RIN4 is a negative regulator of PTI and acts a molecular “phosphoswitch” to control callose deposition and stomatal closure in response to pathogen perception (Ray et al., 2019). Because of its role in PTI, RIN4 is the target of many *P. syringae* effectors and is guarded, in turn, by a number of resistance proteins in a number of different plants: RPM1, RPS2, Ptr1, Mr5, and Rpa1 (Mackey et al., 2003; Mackey et al., 2002; Mazo-Molina et al., 2020; Vogt et al., 2013; Yoon and Rikkerink, 2020). Psa3 effectors AvrRpm1a and HopZ5a have been shown to target RIN4 (Choi et al., 2021; Yoon and Rikkerink, 2020). As RIN4 is evolutionarily conserved across monocot and dicot crops, with promising homologs identified in *Actinidia*, resistance proteins guarding RIN4 and its associated proteins could be durable targets for resistance breeding, with potentially broad-spectrum recognition that could be deployed in a range of cultivated *Actinidia* spp.

The pathogenicity assays in this study of *A. chinensis* var. *chinensis* ‘Hort16A’ and *A. arguta* AA07_03 are among the first to test the virulence of all five described Psa biovars. Psa5 has previously been identified as weakly virulent in the field, while Psa6 has an unknown degree of pathogenicity (Fujikawa and Sawada, 2016; Sawada and Fujikawa, 2019). Similar to Psa3, Psa6 appears to be highly pathogenic in ‘Hort16A’ but avirulent in AA07_03. We confirmed Psa5 as being less pathogenic in ‘Hort16A’, similar to the pathogenicities of Psa1 and Psa2 (Figure 9). The strain-specific level of resistance in AA07_03 across the different Psa biovars suggests that there is a complex resistance gene/avirulence effector relationship present (Salgon et al., 2017). The only partially increased virulence of Psa1 J-35 and Psa2 K-28, relative to that of Psa3 V-13, suggests that these strains may still carry effectors that trigger resistance in AA07_03, including those shared with Psa3 V-13 (Figure 9). Notably, Psa1 carries *avrRpm1a* while Psa2 carries *hopF1c* (and *avrRpm1c*), but Psa 1 and Psa2 may possess other effectors that suppress ETI for these effectors. Nevertheless, the effector presence/absence analysis between these biovars of Psa suggests a hierarchy of recognition strengths in AA07_03. Namely, HopAW1a recognition confers the strongest growth restriction; Psa1 and Psa2 lacking this effector (as well as *hopZ5a*) have the most growth in AA07_03. HopZ5a/AvrRpm1a/HopF1c confer a similar, lower degree of quantifiable resistance, with effector interplay playing a complex role.

The four avirulence effectors that trigger resistance in AA07_03 can be used to identify cognate resistance proteins and can contribute to effector-assisted breeding in kiwifruit cultivar development programmes. Resistance genes that target “Achilles’ heel” effectors which are conserved across epidemic strains and biovars may confer durable, broad-spectrum resistance (Vleeshouwers and Oliver, 2014). For example, *avrRpm1a* is present in Psa1, Psa3 and Psa6, and the closely related *avrRpm1c* is present in Psa2 and Psa5. If these effectors are recognized by the same resistance gene, this might represent a true Achilles’ heel for the whole Psa pathovar. Interestingly, testing of the AvrRpm1c allele from Psa2 K-28 suggested that it is also recognized by AA07_03, possibly by the same resistance protein recognizing AvrRpm1a (Figure S12). Resistance proteins that target effectors that are variable between strains or biovars are of lower priority for resistance breeding, as they are effective only against a subset of the pathogen population. Unfortunately, *hopZ5a* is unique to the pandemic lineage of Psa3. Similarly, *hopF1c* is absent from Psa1 and Psa6, and *hopAW1a* is absent in Psa1 and Psa2. Of further concern around the utility of resistance against EEL-based effectors, genes located upon the same element could easily be inactivated as a block in a single genetic event, as predicted by Rikkerink et al. (2015). This has already been observed in the field isolate Psa3 X-27, with the deletion of five EEL effectors. This highlights the potential for effector loss under selection pressure from resistant plants in the field. This field-based adaptation underscores the importance of deploying durable resistance genes that ideally target conserved effectors with a virulence requirement, which would impose a fitness cost to a pathogen attempting to escape host recognition.

In contrast, the sequential multiple-effector knockout strategy did not show an additive increase in pathogenicity of Psa3 V-13 in AA07_03. In fact, the quadruple avirulence-effector knockout strain (*ΔhopAW1a*/*ΔhopF1c*/*ΔavrRpm1a*/*ΔhopZ5a*) also had reduced pathogenicity in susceptible *A. chinensis* var. *chinensis* ‘Hort16A’ plants. In addition, the increased pathogenicity of the different Psa biovars in *A. arguta* reflected reduced pathogenicity in *A. chinensis* var. *chinensis*, suggesting a trade-off present in the effector repertoire of Psa. This may be a reason for the particularly virulent disease reported during the pandemic spread of Psa3, but not of Psa1 and Psa2, which were earlier emergent diseases of kiwifruit (McCann et al., 2017; Sawada and Fujikawa, 2019). Here it is important to point out that these earlier outbreaks occurred in Korea and Japan where the indigenous *Actinidia* species include *A. arguta* and these biovars therefore presumably evolved partly in the wild *Actinidia* germplasm in Korea/Japan. Taken together, these findings highlight a second route to durable resistance: stacking resistance recognition in plants whereby evasion of resistance through loss of multiple effectors will result in cumulative reduced fitness in the plant host.

Breeding resistance genes into targeted kiwifruit cultivars is essential for long-term management of Psa. Moreover, breeding *durable* resistance requires an understanding of which pathogen effectors are required for virulence and which trigger resistance in potential hosts. The optimal situation is one where resistance genes target essential effectors, as the loss of an essential effector reduces pathogen fitness *in planta*. Loss of these effectors is, therefore, likely to be selected against. Once identified, resistance genes can be introduced into crops. Traditional breeding can be time-consuming and slow new cultivar development (Kim and Kim, 2019). Alternatively, modern GM technology can efficiently introduce resistance genes without linkage drag of undesirable agronomic traits, to create elite transgenic cultivars (Jayaraman et al., 2016). Transgenic crops can also be used to confirm the efficacy of resistance genes before traditional crosses enter pre-commercial field trials, speeding up the cultivar development pipeline. Future research will entail characterizing avirulence effector function, interplay and redundancy to identify which resistance genes are durable breeding targets. Introducing durable Psa resistance that will be effective against the broad spectrum of Psa biovars into future *Actinidia* cultivars will reduce the burden of disease on the horticultural economy and allow a shift towards sustainable production.

## Experimental Procedures

### Leaf tissue immunolabelling & microscopy

Pieces of *A. chinensis* var. *chinensis* ‘Hort16A’ or *A. arguta* AA29_01 leaf, spray-inoculated with Psa3 ICMP 18884 at 10^8^ cfu/mL and harvested at 1–5 days post-infection (dpi), were fixed in 2% paraformaldehyde and 0.1% glutaraldehyde in 0.1M phosphate buffer at pH 7.2 for 1 h under vacuum. Tissue was washed in buffer three times, dehydrated in an ethanol series and embedded in LR White resin (London Resin, Reading, UK) (Sutherland et al., 2009). Sections, 200 nm thick, were cut and dried onto Poly-L-Lysine-coated slides, and left overnight on a hot plate at 45–50°C. These sections were then immunolabelled (Miles et al., 2009; Rheinländer et al., 2013; Sutherland et al., 2009). Briefly, sections were rinsed in Phosphate-Buffered Saline/Tween® (PBS-T), blocked with 0.1% (w/v) bovine serum albumin (Bsa-c, Aurion, Wageningen) in PBS-T for 15 min, and incubated in anti-(1→3)-β-D-glucan antibody (BioSupply, Parkville, Australia) diluted 1:100 in blocking buffer overnight at 4°C. Sections were then washed in PBS-T, incubated for 1 h in Alexa Fluor® 488 goat anti-mouse antibody (Molecular Probes, Eugene, Oreg., USA) diluted 1:600 in PBS, washed in PBS-T, followed by ultrapure water and mounted in Citifluor (Leicester, UK). Sections were viewed on an Olympus Vanox AHTB3 microscope using an interference blue excitation filter set and images collected with a Roper Scientific CoolSnap color digital camera. To highlight the leaf cell walls, sections were either stained with 0.01% (w/v) calcofluor in water (labeling cellulose) or immunolabelled with LM19 (labeling pectin) in a process that followed the initial labelling. The immunolabeling protocol was similar to that described above except that Alexa Fluor® 594 goat anti-rat (Molecular Probes) was used as the secondary antibody/fluorchrome combination. The hypersensitive response (HR) was observed by destaining the tissue in acetic acid:ethanol (1:3) for 8 h, washed in 100% ethanol and observed in bright field through the Olympus microscope.

### Field survey & Psa isolation

Samples were taken from leaf spots on vines in the Plant & Food Research Te Puke Research Orchard *Actinidia* germplasm collection. Infected leaves, fruit, bud, shoot and cane samples were taken using secateurs sterilized with 80% ethanol. A 1-cm diameter cork borer was used to punch three leaf discs from each symptomatic leaf. Leaf discs were surface-sterilized in 70% ethanol for 10 s, and washed with sterile MilliQ water in a Falcon tube. For each sample, three leaf discs were placed into an Eppendorf™ Microcentrifuge Safe-Lock™ tube (Fisher Scientific, California, United States) with 350 µL sterile 10 mM MgSO_4_ and three sterile 3.5-mm stainless steel beads. Samples were ground for two runs of 1 min at the maximum speed in a Storm24 Bullet Blender (Next Advance, New York, United States). Tubes were vigorously inverted to resuspend the leaf material pellets between each grinding run. Supernatant (200 µL) was then spread onto lysogeny broth (LB) agar plates (Bertani, 1951) supplemented with 12.5 µg/mL nitrofurantoin and 40 µg/mL cephalexin and incubated for 48 h at 22°C. The bacterial lawn was then re-streaked onto new LB agar plates (supplemented with nitrofurantoin and cephalexin) until single colonies could be isolated.

Quantitative PCR (qPCR) was carried out on an Illumina Eco Real-Time PCR platform (Illumina, Melbourne, Australia), following the protocol outlined by Barrett-Manako and colleagues (Barrett-Manako et al., 2021). Single colonies were tested with Psa-*ITS*, Psa HopZ5-F2/R2 and HopA1-F2/R1 qPCR primers to identify Psa3 strains (Andersen et al., 2017; Table S1). Samples that amplified in under 35 qPCR cycles were prepared as a 20% (w/v) glycerol stock for long-term storage.

### DNA extraction & sequencing

DNA was purified following the Gentra® Puregene® protocol for Gram-negative bacteria (Qiagen, Hilden, Germany). Libraries were constructed using the Nextera DNA preparation kit and sequenced on an Illumina Hi-Seq 2500 platform (paired-end 125 bp reads) (Illumina). Quality control reports for the raw sequencing reads were generated using FastQC (Andrews, 2010). Raw sequencing reads underwent quality and adapter trimming using BBDuk (Bushnell, 2014) (version 38.62; parameters: ktrim=r, k=2,1 mink=11, hdist=2, xminlen=50, ftm=5, tpe, tbo, qtrim=r, trimq=10, minlen=50, maq=10). Trimmed reads were mapped to the reference genome Psa ICMP 18884 using the bwa aligner (Li and Durbin, 2010) and variants were called using bcftools (Li, 2011) (version 1.9). Bedtools genomecov was used to generate .bed files of regions with low or no coverage (Quinlan and Hall, 2010). Bcftools was then used to generate a consensus sequence, masking regions of low or no coverage (Li, 2011). Reference genome sequences for the Psa strains used in this study (Templeton et al., 2015; Table 2) were obtained from the NCBI GenBank. All downstream analyses were carried out in Geneious (Kearse et al., 2012; version 10.0.9).

**Table 2.**
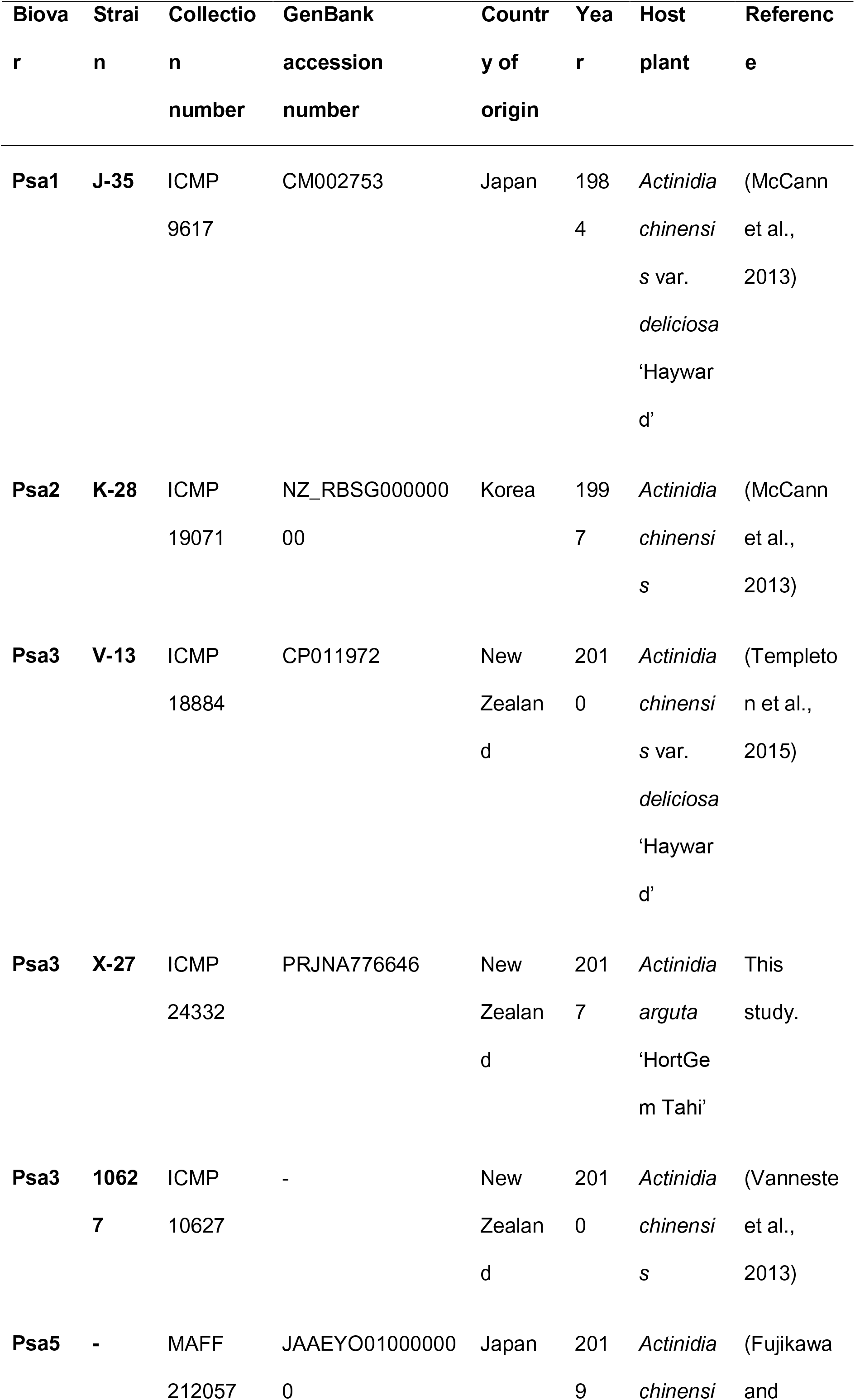
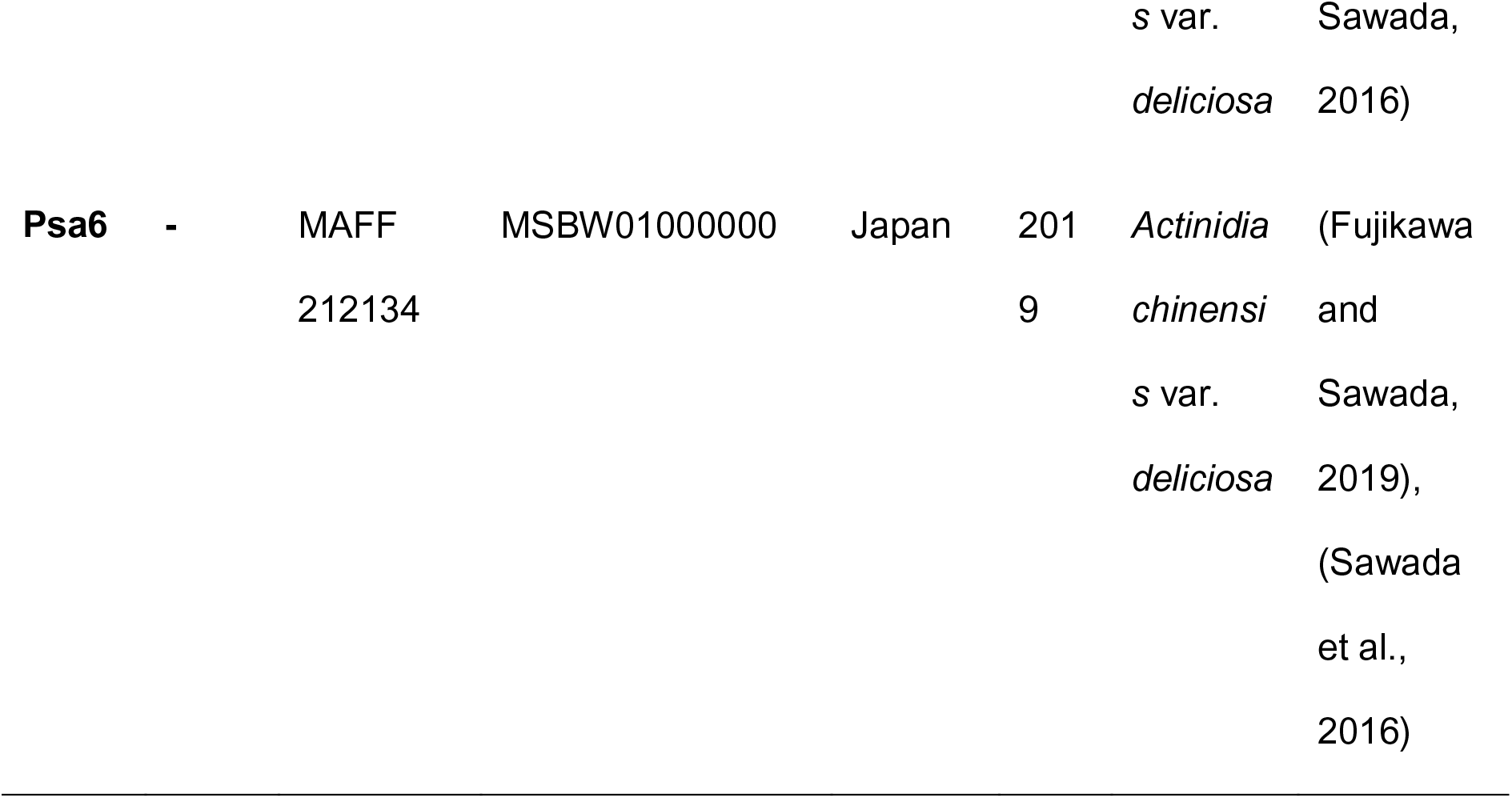
Wild-type Psa strains. All wild-type Psa strains were sourced from ICMP/MAFF.

### Microbiological methods

Psa strains used in this study are listed in Table 1 and Table 2. All Psa strains were streaked from glycerol stocks onto LB agar supplemented with appropriate antibiotics; plates were sealed and grown for 48 h at 22°C. Overnight shaking cultures were grown in LB supplemented with appropriate antibiotics and incubated at 22°C with 200 rpm shaking. LB agar was supplemented with 12.5 µg/mL nitrofurantoin (Sigma Aldrich, New Zealand) and 40 µg/mL cephalexin (Sigma Aldrich) for Psa selection. To select for Psa strains carrying pK18mobsacB, LB agar was supplemented with 50 µg/mL kanamycin. To counter-select against Psa strains carrying pK18mobsacB, LB agar was supplemented with 12.5 µg/mL nitrofurantoin, 40 µg/mL cephalexin, and 5% sucrose (Merck Millipore, New Zealand). To select for Psa strains carrying pBBR1MCS-5B vectors for effector complementation, LB agar was supplemented with 50 µg/mL gentamicin (Sigma Aldrich).

### Rooted plant inoculations and testing

Experiments were conducted as described previously in Vanneste et al. (2013). Briefly, a bacterial suspension for Psa3 X-27 or Psa3 ICMP 10627 (WT; clonal isolate related to Psa3 ICMP 18884; Vanneste et al., 2013) was made in water from freshly grown colonies on King’s B agar plates (King et al., 1954) and adjusted to ∼10^8^ cfu/mL. Suspensions were sprayed onto the abaxial side of all leaves of three 3- to 4-month-old seedlings of *A. arguta* AA07_03 or *A. chinensis* var. *chinensis* ‘Hort16A’. Plants were kept at approximately 20°C in plastic chambers to maintain the relative humidity. Leaf samples were taken at 14 dpi to re-isolate bacterial DNA for PCR confirmation using *Psa-ITS* and *Psa-ompP1* primers (Table S1) as described previously (Vanneste et al., 2013). Leaf symptomology photographs were taken at 6 months post-infection.

### Psa3 effector gene knock-out library

Psa3 ICMP 18884 (hereafter referred to as Psa3 V-13) was used as the WT for a Psa effector knockout library using the pK18mobsacB-based system. A complete library of 25 Psa3 V-13 effector knockout strains was developed with effectors knocked out either individually, in pairs if homologs were present (*hopAM1a-1*/*hopAM1a-2*) or as a functional group (CEL, EEL various iterations, *hopZ5a*/*hopH1a*, or *hopQ1a*/*hopD1a*) (Table 2). Effector knockout plasmids were developed for Psa3 V-13 using the methodology established by Kvitko and Collmer (2011) and as described in Jayaraman et al. (2020). Briefly, flanking regions 1kb upstream (UP) and 1kb downstream (DN) of the effectors of interest were PCR-amplified with UP-R and DN-F cloning primers carrying an inserted *Xba*I site (Table S1), digested with *Xba*I restriction enzyme (New England Biolabs/NEB, MA, USA), and ligated to form a 2 kb knockout fragment. This 2 kb fragment was subsequently cloned into the *Eco53k*I restriction enzyme (NEB) site of pK18B-E (Jayaraman et al., 2020). The knockout fragment sequence and quality were verified by sequencing using M13F and M13R primers (Macrogen, South Korea). Psa3 V-13 was transformed with each knockout vector by electroporation (see Plasmid transformation section below). Transformants were plated onto LB agar supplemented with kanamycin to select for strains carrying a genomic insertion of the pK18B-E knockout construct. Resultant colonies were streaked onto LB agar supplemented with 5% sucrose to counter-select against the *sacB* gene in pK18B-E. Resulting colonies were then screened using PCR (check-F/R) primers that amplified outside the knockout region (Table S1). Successful knockout strains were sub-cultured from 5% sucrose plates onto LB agar supplemented with or without 50 µg/mL kanamycin to confirm plasmid loss and restored kanamycin sensitivity, and the ∼2 kb knockout fragment PCR amplicon was sequenced to confirm authenticity (Macrogen, South Korea). The Psa3 Δ*CEL* and Psa3 Δ*hopR1* strains included in the effector knockout strain library were described and characterized earlier (Jayaraman et al., 2020).

### Psa3 complete effector knockout

A Psa3 V-13 complete effector knockout strain was generated as for the single knockouts using the same pK18mobsacB-based vectors used before. The single effectors or blocks of effectors were sequentially knocked out to make the 30 effector knockout strain (Psa3 V-13 Δ*30E*) in the order: *hopZ5a*/*hopH1a* (using the *hopZ5a*/*hopH1a* double knockout vector), *hopBP1a* (previously *hopZ3*), *hopQ1a*, *hopAS1b*, *avrPto1b* (previously *avrPto5*), *avrRpm1a*, *fEEL* (*avrD1*/*avrB2b*/*hopF4a*/*hopAW1a*/*hopF1e*/*hopAF1b*/*hopD2a*/*hopF1a*), hopF1c (previously *hopF2*), *hopD1a* (using the *hopQ1a*/*hopD1a* double knockout vector), *CEL* (*hopN1a*/*hopAA1d*/*hopM1f*/*avrE1d*), *hopR1b*, *hopAZ1a*, *hopS2b*, *hopY1b*, *hopAM1a-1*, *hopAM1a-2*, *hopBN1a*, *hopW1c* (previously *hopAE1*), *hopAU1a*, and *hopI1c*. Knockouts were confirmed by PCR. Effectors that did not have a functional type III secretion signal owing to truncation or disruption, or did not possess a HrpL box promoter individually or in an operon (confirmed by expression analysis in McAtee et al. (2018)) were not knocked out and included the following effector loci: *avrRpm1c*, *hopA1a*, *hopAA1b*, *hopAG1f*, *hopAH1c*, *hopAI1b*, *hopAT1e* (previously *hopAV1*), *hopAB1b* (previously *hopAY1*). The effector *hopAA1d* was also considered a pseudogene under these criteria but was knocked out with other effectors in the *CEL*.

### Avirulence effector cloning

Psa3 V-13 avirulence effector genes *hopAW1a*, *hopZ5a*, *avrRpm1a*, or *hopF1c* (along with its chaperone *shcF*) were PCR-amplified using Q5 High fidelity polymerase (NEB) from Psa3 V-13 genomic DNA using cloning primers (Table S1) including their HrpL box promoters. PCR amplicons were gel-extracted from agarose, and cloned by blunt-end ligation into the *Eco53k*I restriction enzyme site in the pBBR1MCS-5 broad host range vector. Clones were confirmed by sequencing (Macrogen).

### Plasmid transformations into Psa3

Effector genes were plasmid-complemented back into Psa3 V-13 Δ*sEEL* or Psa3 V-13 Δ*30E* following methodology established in Jayaraman et al. (2020). Psa strains were inoculated into 5 mL LB supplemented with appropriate antibiotics and incubated overnight at 20°C until mid-log phase was reached (3×10^8^ cfu/mL). Cultures (2 mL) were collected by centrifugation at 17,000 *g* at 4°C and washed in cold sterile water multiple times to induce electro-competency according to the previously defined protocol (Choi et al., 2006). The final bacterial pellets were resuspended in 100 µL sterile 300 mM sucrose solution, and plasmid DNA added (200–500 ng per reaction). Electro-competent Psa cells were transformed on the Gene Pulser Xcell™ Electroporation System (Bio-Rad, New Zealand), supplemented with sterile, antibiotic-free LB and incubated at 22°C for 1 h with 200 rpm shaking, before plating onto LB agar supplemented with gentamicin for plasmid selection and incubated for 48–96 h at 22°C.

### Pathogenicity assays

*Actinidia* spp. plantlets were obtained from Multiflora Laboratories (Auckland, New Zealand). Plants were grown in 400-mL lidded plastic ‘pottles’ on half-strength Murashige and Skoog (MS) Agar, with 3–5 plantlets per pottle. Plantlets were grown in a climate-controlled room at 20°C with a 16 h/8 h light/dark cycle and used within 2–3 months. Plantlets were infected using an *in planta* flooding assay, as established in McAtee et al. (2018). Briefly, kiwifruit plantlets were inoculated by flooding with 500 mL Psa inoculum (∼5 x 10^6^ cfu/mL) for 3 min, and grown in a climate room at 20°C with a 16 h/8 h light/dark cycle. Un-inoculated plantlets were occasionally checked throughout the experiments for Psa contamination and none was detected.

To quantify bacterial growth of Psa *in planta*, leaf samples were taken at 6 or 12 dpi. A 0.8-cm diameter cork borer was used to punch four leaf discs per replicate, with four pseudobiological replicates taken per pottle (n = 16), surface-sterilized, and each ground in 350 µL sterile 10 mM MgSO_4_ with three 3.5-mm stainless steel beads using a Storm24 Bullet Blender (Next Advance, NY, USA). Leaf homogenate stored at −20°C overnight prior to PDQeX DNA extraction according to a previously described protocol (Jayaraman et al., 2021).

A serial dilution of leaf homogenate was prepared to quantify cfu/cm^2^ by the plate count method. A 10-fold dilution series of leaf homogenate in sterile 10 mM MgSO_4_ was made, to a final dilution of 10^-5^ (*A. arguta*) or 10^-7^ (*A. chinensis)*. Each 10-fold dilution in the dilution series was spot-plated (10 µL) onto LB agar supplemented with appropriate antibiotics. Plates were incubated for 48 h at 20°C and resultant colonies were counted to calculate the cfu/cm^2^. To assess disease phenotypes, plantlets were inoculated at ∼1 x 10^8^ cfu/mL and observed at 50 dpi as established by Jayaraman and colleagues (Jayaraman et al., 2021). A modified PIDIQ Image-J macro script (Laflamme et al., 2016) was used to assess leaf yellowing and browning.

### Quantitative PCR

Real-time quantitative PCR (qPCR) was carried out on an Illumina Eco Real-Time PCR platform, following the protocol outlined in Barrett-Manako et al. (2021), with the annealing temperature lowered to 57°C to improve the efficiency of the *EF1α* SN126 L/R primers. The primers used for qPCR are listed in Table S1.

### Ion leakage

Psa3 V-13 Δ*30E* (Table 1) or *P. fluorescens* (T3S or WT; Table S2) carrying empty vector or effector constructs were streaked from glycerol stocks onto LB agar plates with antibiotic selection, were grown for 2 days at 22°C, and were restreaked on fresh agar media, and were allowed to grow overnight. Bacteria were then harvested from plates, were resuspended in 10 mM MgCl_2_, and were diluted to ∼10^8^ cfu/mL. Vacuum-infiltrations were carried out using a pump and glass bell. Leaves were harvested from the tissue culture tubs and were submerged in 30 ml of bacterial inoculum. The vacuum was run until bubbles were rapidly forming. The vacuum valve was then shut and the air slowly let back in. The infiltration was repeated a second time for those leaves not fully infiltrated and any remaining non-infiltrated leaves were removed, as determined by visual examination. For each treatment, leaf discs (6 mm diameter) were harvested from the uniformly vacuum-infiltrated leaf area and were washed in distilled water for 1 h. Six discs were placed in 3 ml of water, and conductivity was measured over 48 h, using a LAQUAtwin EC-33 conductivity meter (Horiba). The standard errors of the means were calculated from five pseudobiological replicates. Data for each timepoint was analyzed by ANOVA followed by a Tukey’s HSD post hoc test.

### Reporter eclipse

Freshly expanded leaves of *A. arguta* AA07_03 were co-bombarded with DNA-coated gold particles carrying pRT99-GUS and pICH86988 with the effector of interest, as described in Jayaraman et al. (2021).

### Statistical analysis

Statistical analysis was conducted in R (R Core Team, 2018), and figures were produced using the packages “ggplot2”(Wickham, 2016) and “ggpubr” (Kassambara, 2017). *Post hoc* statistical tests were conducted using the “ggpubr” and “agricolae” packages (de Mendiburu, 2017; Kassambara, 2017). The stats_compare_means() function from the “ggpubr” package was used to calculate omnibus one-way analysis of variance (ANOVA) statistics to identify statistically significant differences across all treatment groups (Kassambara, 2017). For normally distributed populations, Student’s *t*-test was used to conduct pair-wise, parametric *t*-tests between an indicated strain and a designated reference strain (Kassambara, 2017). The HSD.test() function from the “agricolae” package was used to calculate Tukey’s Honest Significant Difference (de Mendiburu, 2017).

## Author Contributions

J.J., C.B., and M.D.T. conceived the work and planned experiments; L.M.H., J.J., P.W.S., M.M., S.A., A.C., R.C., M.A., C.H.M., O. L., M.M.S., J.L.V., and C.B. performed experiments; L.M.H., J.J., C.B., and M.D.T. analyzed data; L.M.H., J.J., and M.D.T. wrote the paper.

## Acknowledgments

We would like to thank Dr Jo Bowen (PFR), Dr Erik Rikkerink (PFR), and Assoc. Prof. Andrew Allan (PFR) for critical reading of this manuscript. This work was funded (including a post-doctoral fellowship to JJ) by the Bio-protection Research Centre (Tertiary Education Commission) and a Rutherford Foundation Post-doctoral fellowship. LMH would like to thank Zespri International for an MSc scholarship.

## Supplementary tables

**Table S1.**
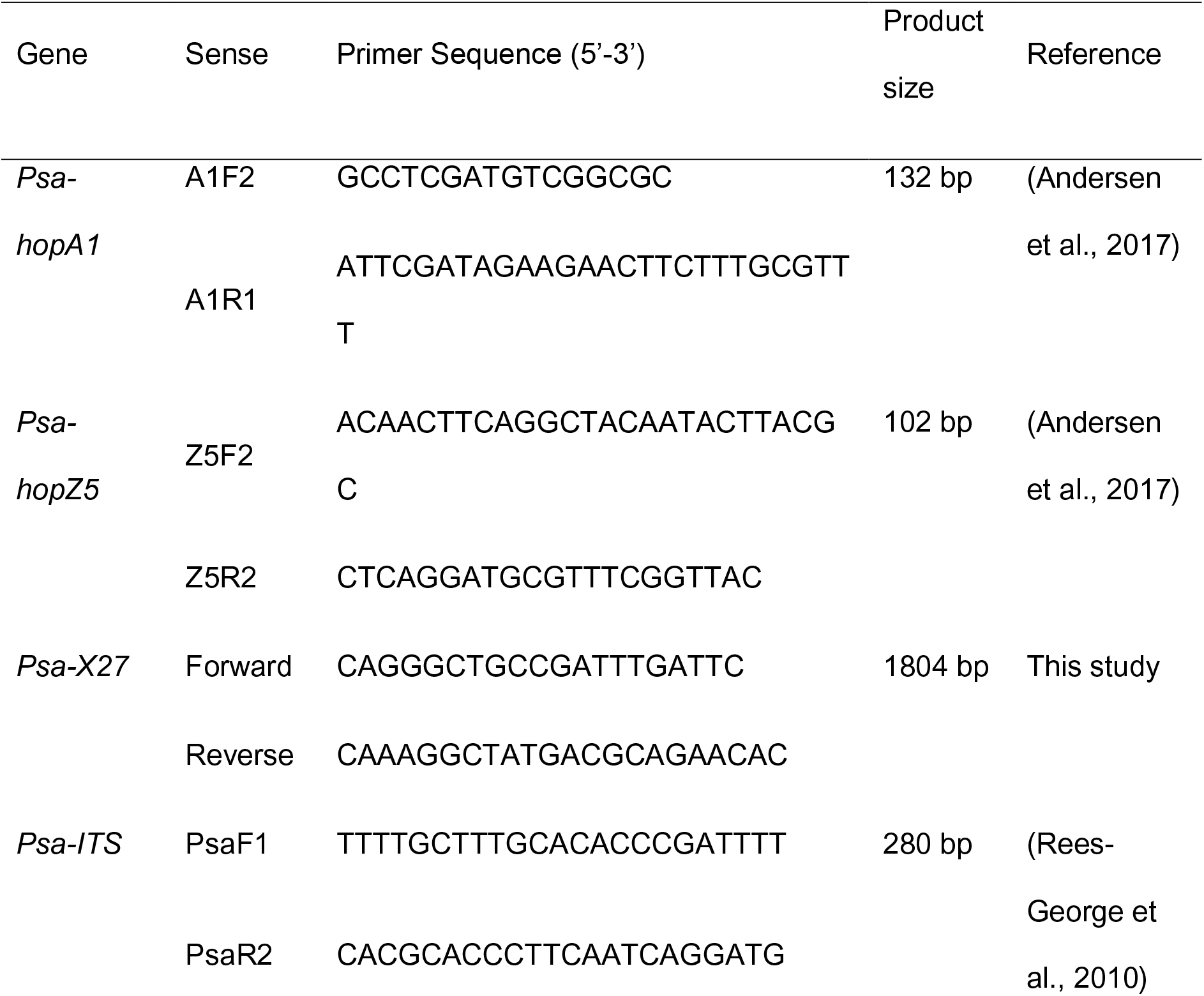
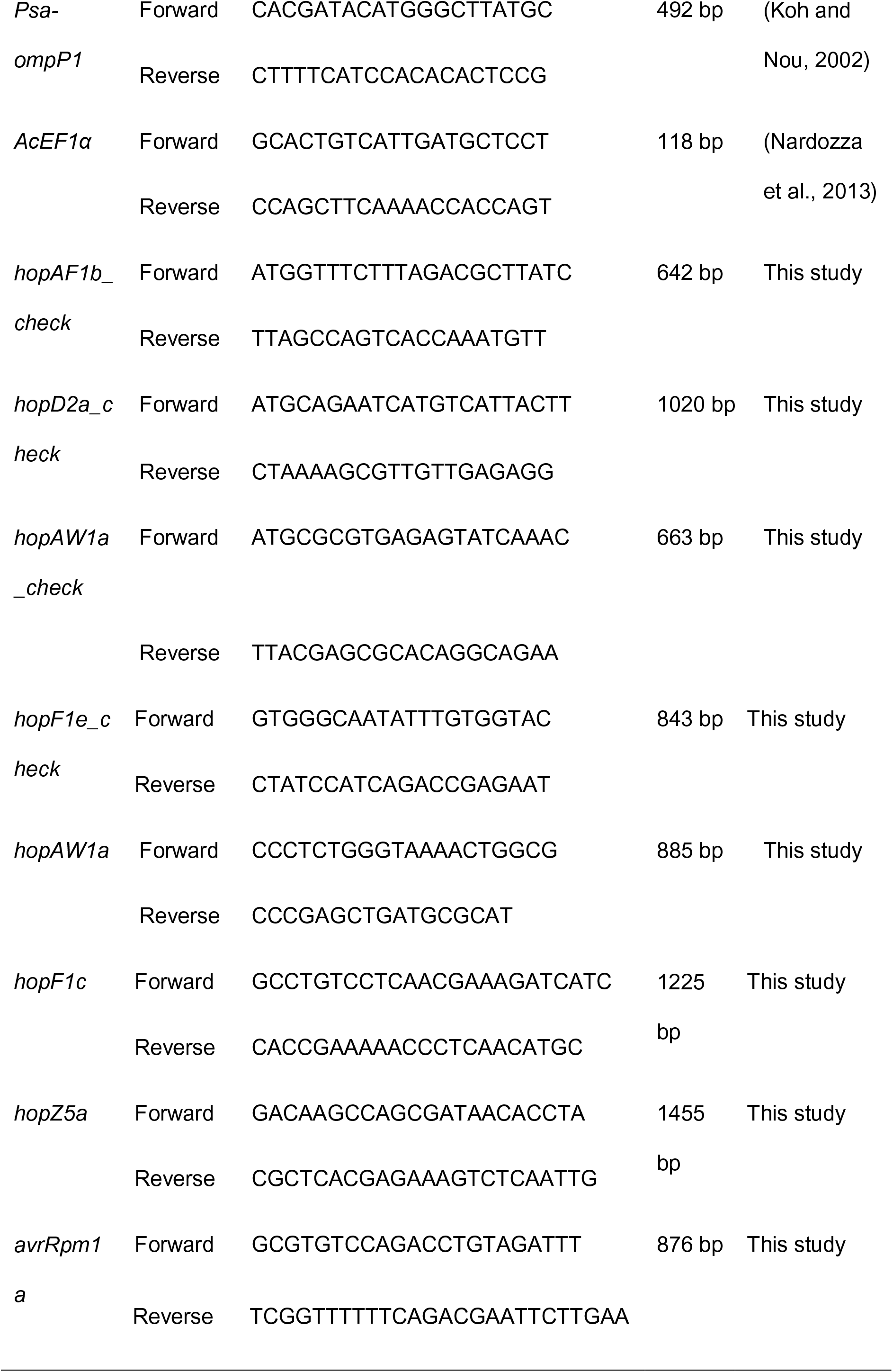
Plasmid cloning and confirmation primers used in this study.

**Table S2.** Pseudomonas fluorescens plasmid-complemented strains used in this study.

| Strain | Description | Source |
| --- | --- | --- |
| *Pfo* (T3S) | *P. fluorescens* Pf0-1 carrying an artificial type III secretion system from *P. syringae* pv. *syringae* 61 | (Thomas et al., 2009) |
| *Pfo* (WT) | *P. fluorescens* Pf0-1 WT strain | (Thomas et al., 2009) |
| *Pfo* (T3S) or *Pfo* (WT) + EV | Plasmid-complemented with empty vector (pBBR1MCS-5) | (Jayaraman et al., 2020) |
| *Pfo* (T3S) or *Pfo* (WT) + hopAW1a | Plasmid-complemented with *hopAW1a* (cloned under *avrRps4* promoter and tagged with HA) | This study |
| *Pfo* (T3S) or *Pfo* (WT) + hopZ5a | Plasmid-complemented with *hopZ5a* (cloned under *avrRps4* promoter and tagged with HA) | This study |
| *Pfo* (T3S) or *Pfo* (WT) + hopF1c | Plasmid-complemented with *hopF1c* (cloned under *avrRps4* promoter and tagged with HA) | This study |
| *Pfo* (T3S) or *Pfo* (WT) +<br>*avrRpm1a* | Plasmid-complemented with<br>*avrRpm1a* (cloned under<br>*avrRps4* promoter and tagged<br>with HA) | This study |
| *Pfo* (T3S) or *Pfo* (WT) +<br>*hopA1j<sub>Psy61</sub>* | Plasmid-complemented with<br>*shcA* and *hopA1j* from *P.*<br>*syringae* pv. *syringae* 61 | (Jayaraman et al., 2021) |

## Supplementary Figures

**Figure S1:**
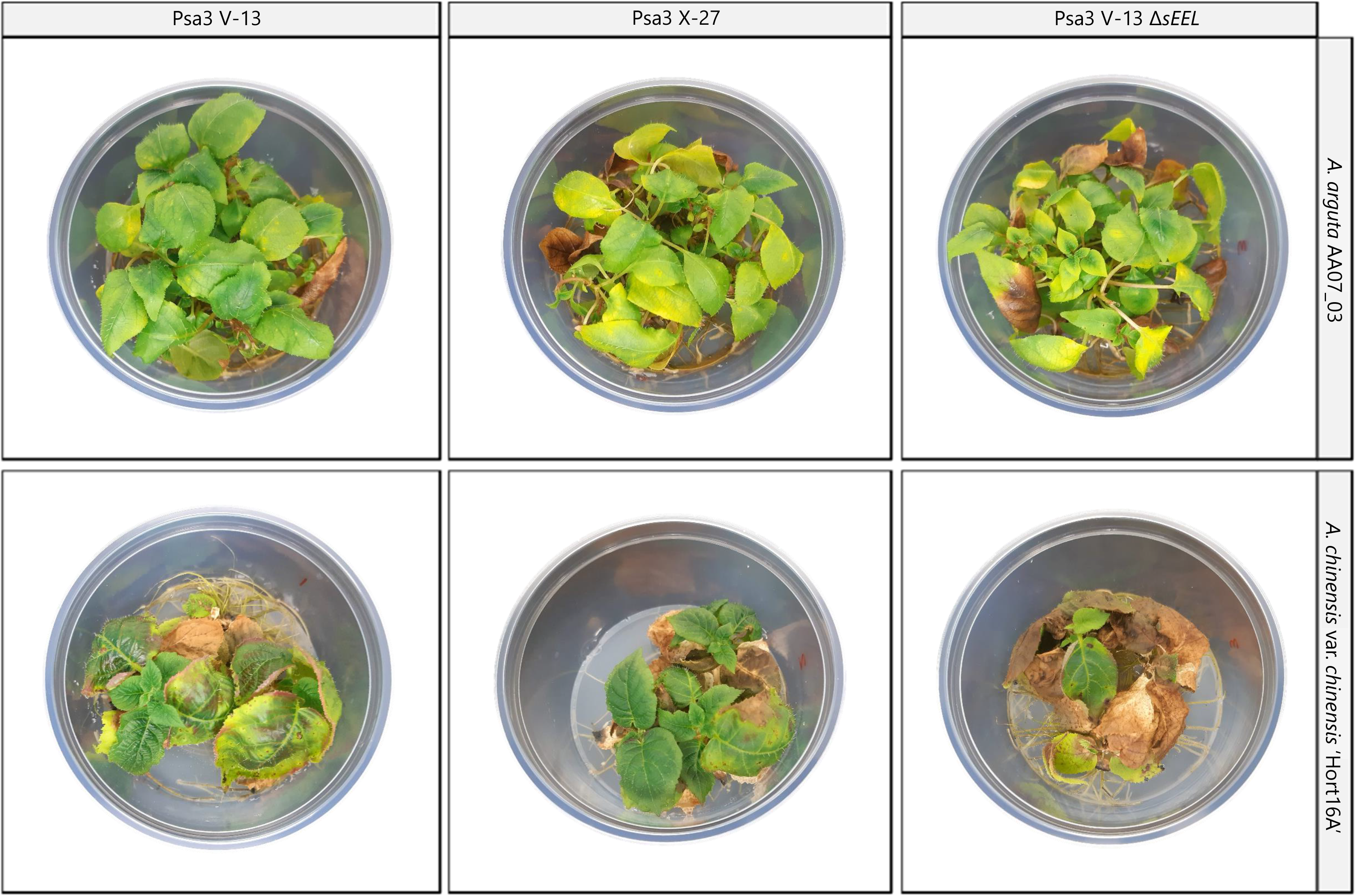
Symptom development of Psa3 V-13, Psa3 X-27, and Psa3 V-13 Δ*sEEL* in *Actinidia arguta* and *A. chinensis* var. *chinensis*. *A. arguta* AA07_03 kiwifruit plantlets were flood-inoculated at approximately 10^7^ cfu/mL. Photographs of symptom development in representative pottles were taken at 50 days post-infection.

**Figure S2:**
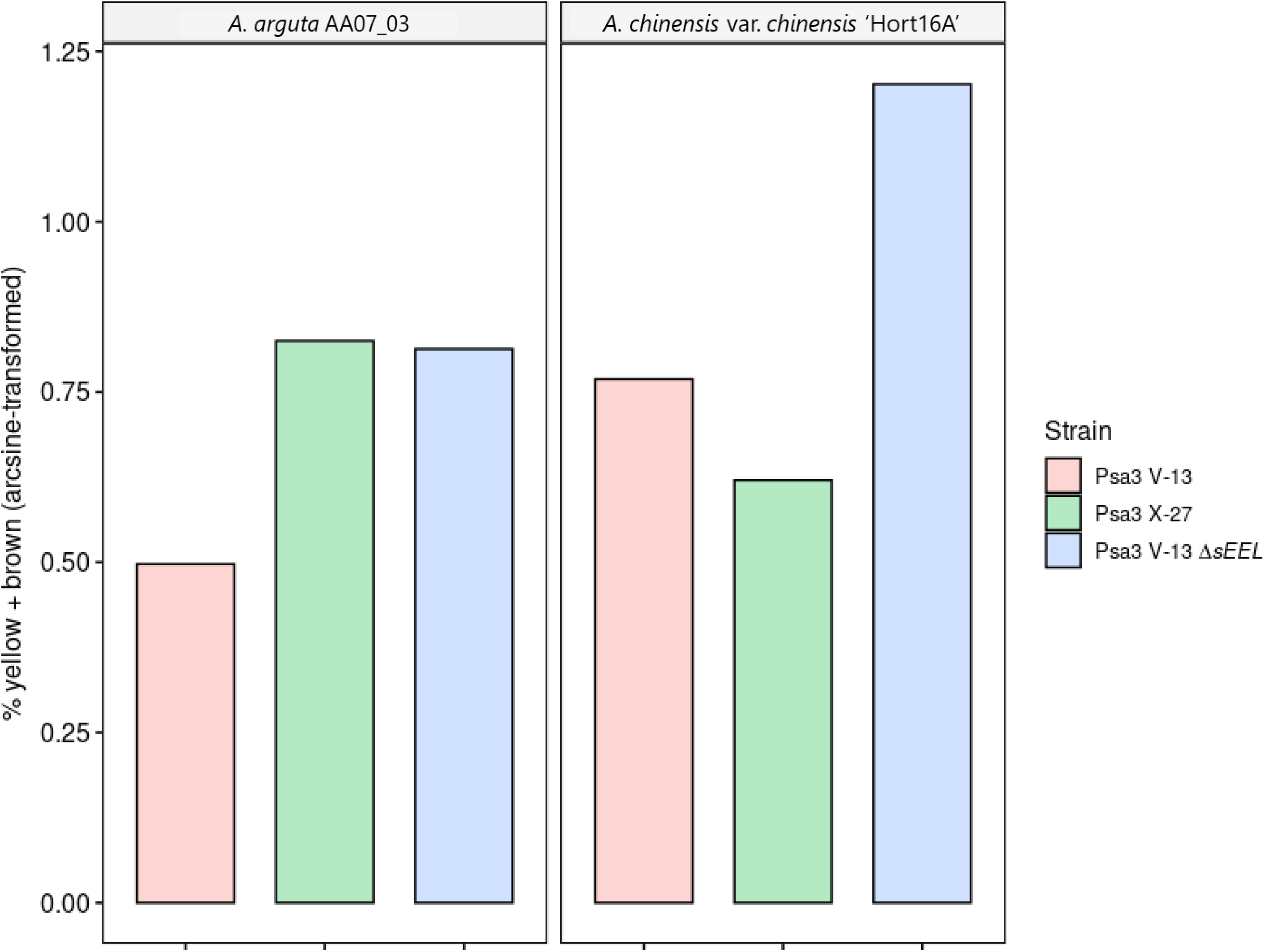
Quantification of symptom development of Psa3 V-13, Psa3 X-27, and Psa3 V-13 Δ*sEEL* in *Actinidia arguta* and *A. chinensis* var. *chinensis*. A modified PIDIQ image-based analysis of leaf yellowing and browning, expressed as a normalized arcsine-transformed percentage for symptomology photographs taken at 50 days post-infection (Figure S1). Methodology adapted and modified from that in Laflamme et al. (2020).

**Figure S3:**
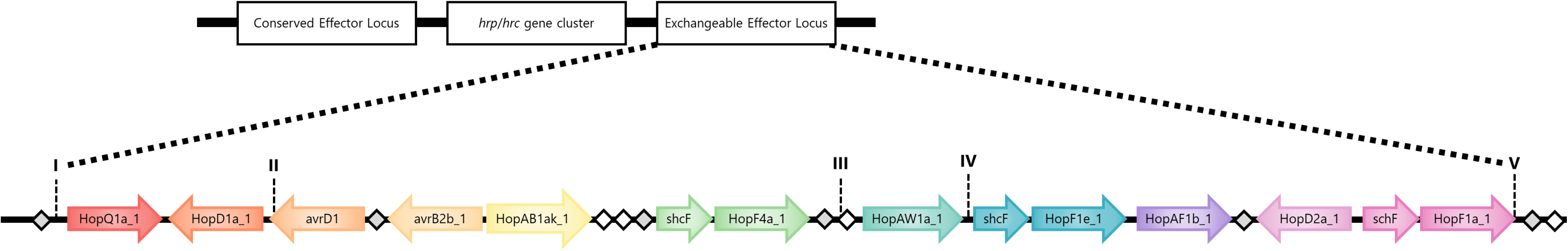
The non-canonical extended exchangeable effector locus (xEEL) encompassing the full EEL (fEEL), short EEL (sEEL), and tiny EEL (tEEL) loci. Schematic of the effectors comprising the xEEL (I-V; *hopQ1a* – *hopF1a*), fEEL (II-V; *avrD1* – *hopF1a*), sEEL (III-V; *hopAW1a* – *hopF1a*), and tEEL (IV-V; *hopF1e* – *hopF1a*) loci in Psa3 V-13 ICMP 18884 strain are indicated. Potential recombination sites are indicated: Miniature Inverted Repeat Transposable Element (MITE; grey diamonds), DDE terminal inverted repeats (black diamonds).

**Figure S4:**
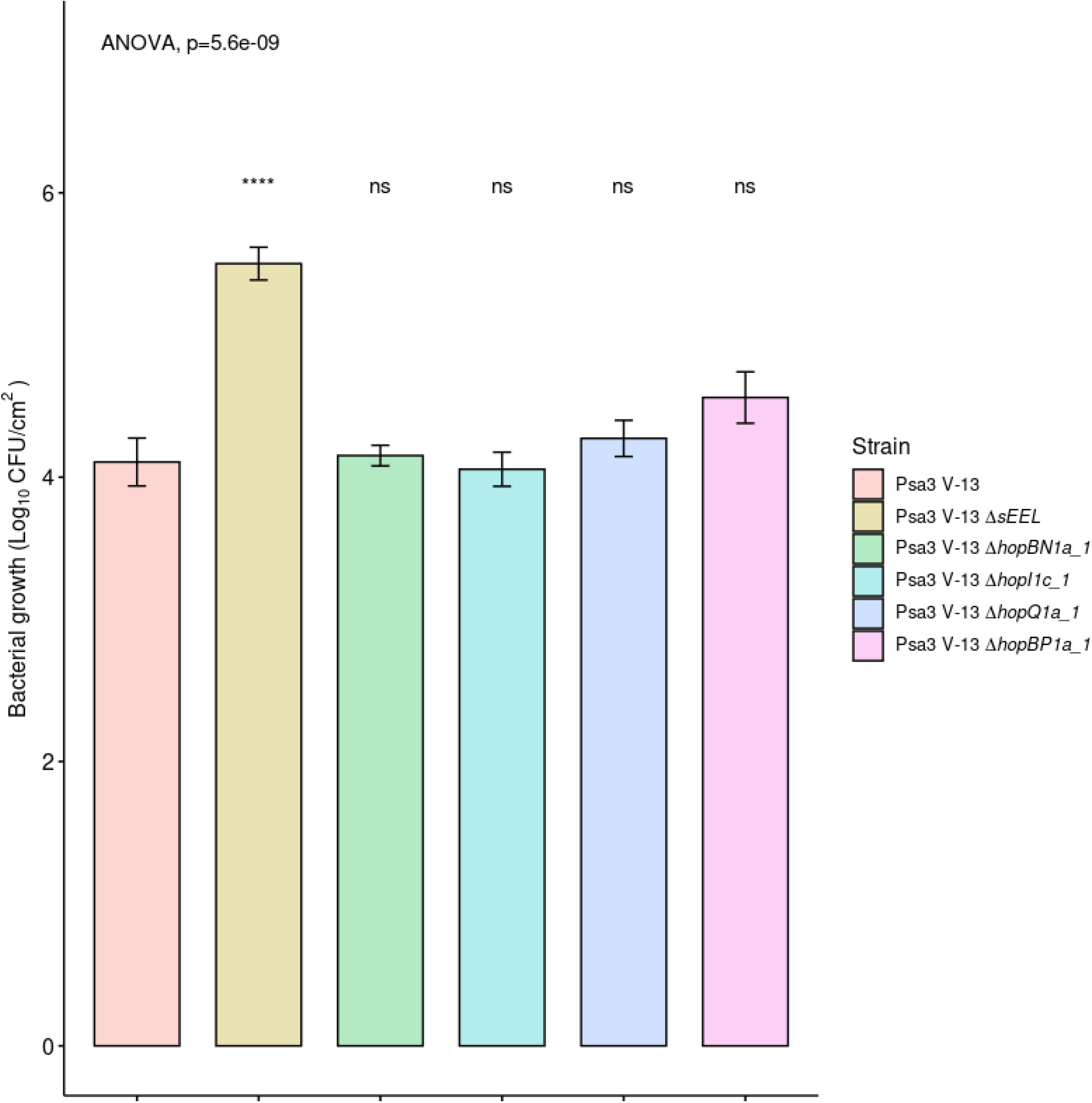
Pathogenicity assay of Psa3 V-13 selected effector knockout strains in *Actinidia arguta* AA07_03 confirming lack of contribution towards avirulence. *A. arguta* AA07_03 kiwifruit plantlets were flood-inoculated at approximately 10^6^ cfu/mL. Bacterial pathogenicity was quantified relative to Psa3 V-13 using plate count quantification for four pseudobiological replicates, per strain, per experimental run and error bars represent the standard error of the mean (SEM). Asterisks indicate the statistically significant difference of Student’s *t*-test between the indicated strain and wild-type Psa3 V-13, where p≤.001 (****), and p>.05 (ns; not significant). This experiment was separately conducted twice (biological replicates) with two batches of independently grown plants and data were stacked to generate the bar graphs shown.

**Figure S5:**
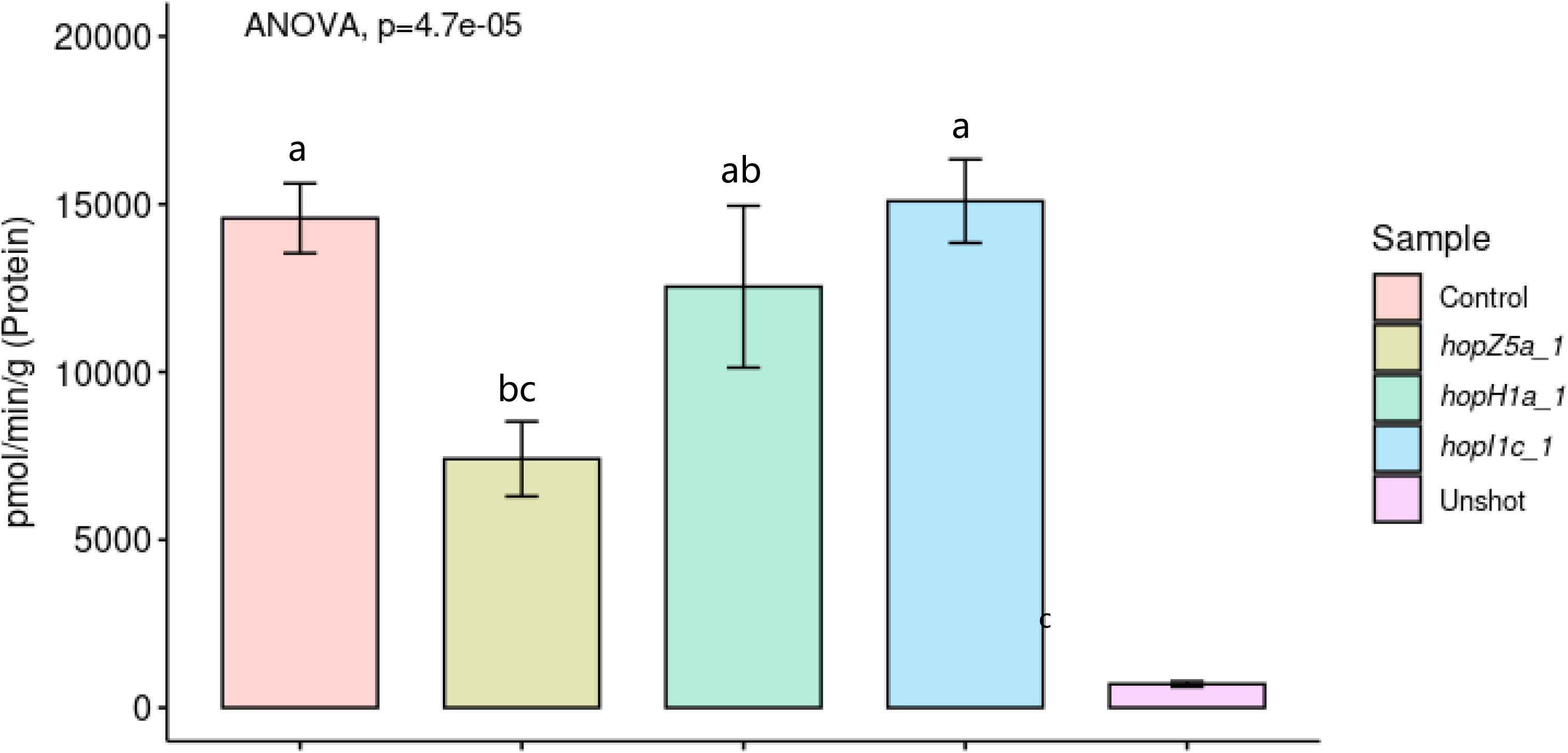
Biolistic transformation reporter eclipse assay demonstrates that HopZ5a, and not HopH1a, triggers a host-specific immunity response in *Actinidia arguta*. Avirulence effectors cloned into binary vector constructs tagged with GFP, or an empty vector (Control), were co-expressed with a β-glucuronidase (GUS) reporter construct using biolistic bombardment and priming in leaves from *A. arguta* AA07_03 plantlets (Jayaraman et al., 2021). The GUS activity was measured 48 hours after DNA bombardment. Error bars represent the standard errors of the means for three independent biological replicates with six technical replicates each (n=18). HopI1c was used as the positive control and un-infiltrated leaf tissue (Unshot) as the negative control. Tukey’s HSD indicates treatment groups that are significantly different at α ≤ 0.1 with different letters.

**Figure S6:**
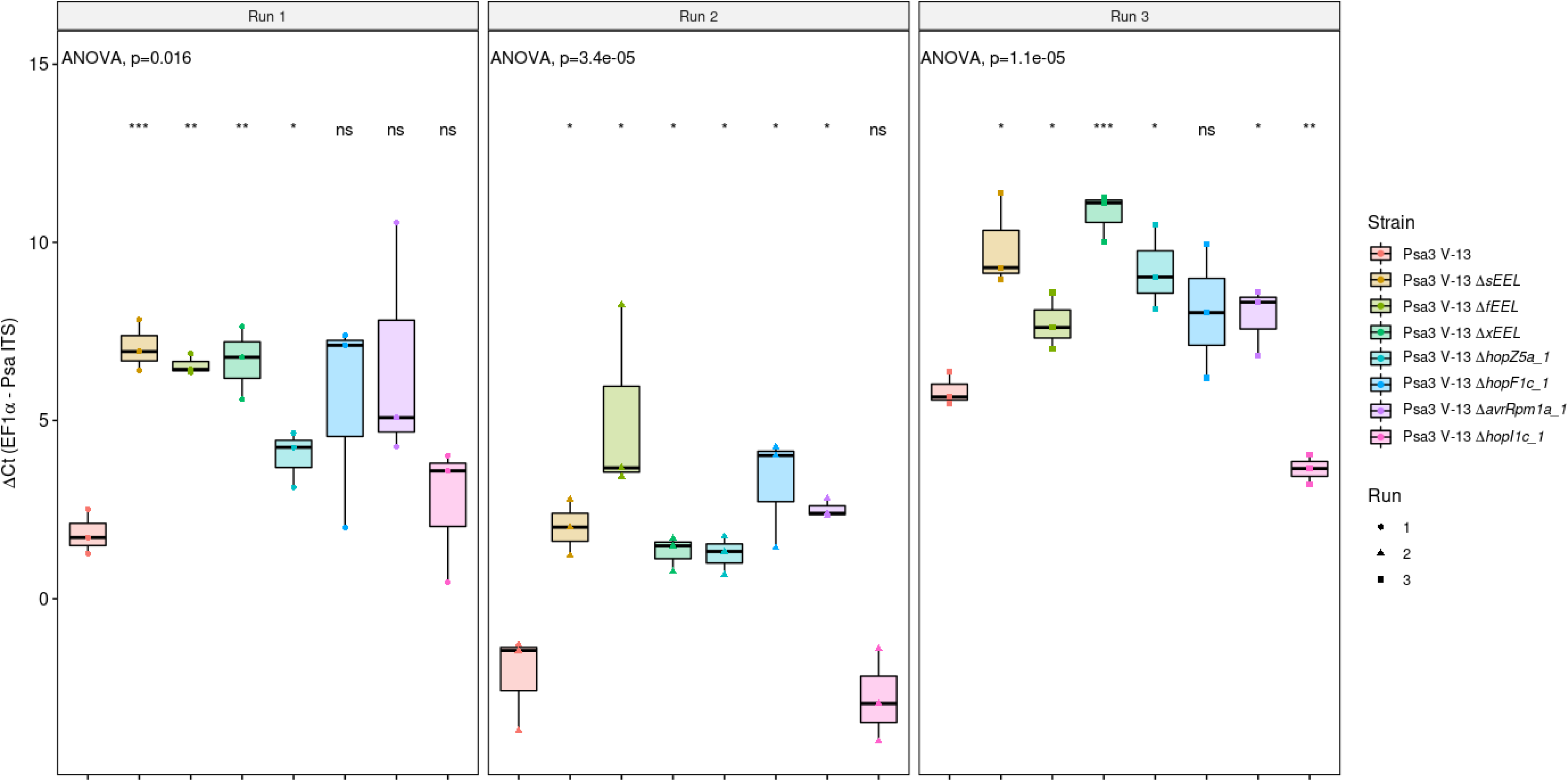
qPCR-based pathogenicity assay of Psa3 V-13 selected effector knockout strains in *Actinidia arguta* confirming recognition of four avirulence loci. *A. arguta* AA07_03 kiwifruit plantlets were flood-inoculated at approximately 10^6^ cfu/mL. Bacterial pathogenicity was quantified relative to Psa3 V-13 using the ΔCt analysis method for four pseudobiological replicates, per strain, per experimental run. Data are presented as box and whisker plots, with black bars representing the median values and whiskers representing the 1.5 inter-quartile range. The data have been faceted by experimental run. Asterisks indicate the statistically significant difference of Student’s *t*-test between the indicated strain and wild-type Psa3 V-13, where p ≤.05 (*), p≤.01 (**), p≤.001 (***), and p>.05 (ns; not significant). These three experiments (biological replications) were separately conducted with three batches of independently grown plants.

**Figure S7:**
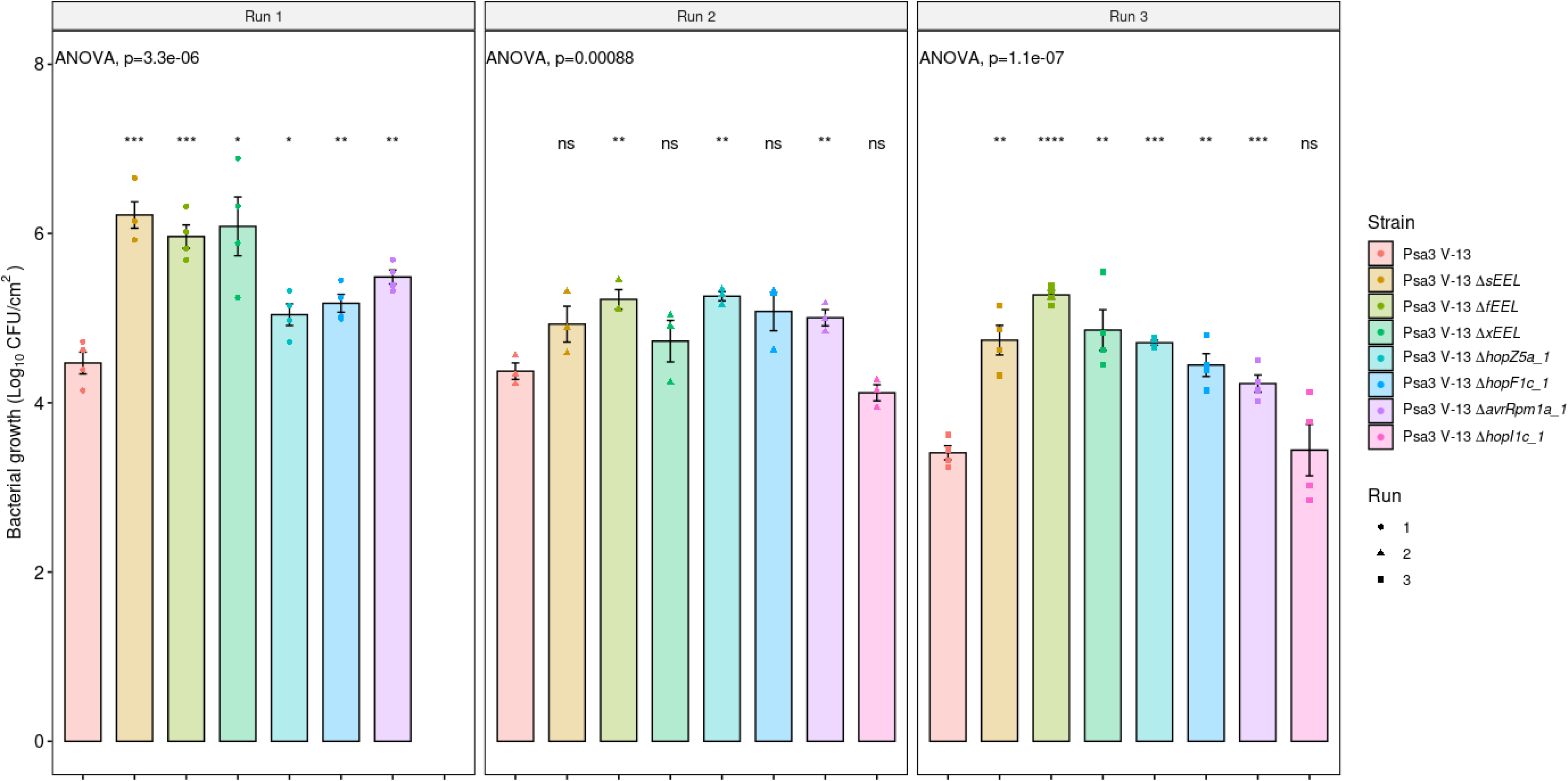
Agarose plate-based pathogenicity assay of Psa3 V-13-selected effector knockout strains in *Actinidia arguta* confirming recognition of four avirulence loci. *A. arguta* AA07_03 kiwifruit plantlets were flood-inoculated at approximately 10^6^ cfu/mL. Bacterial pathogenicity was quantified relative to Psa3 V-13 using plate count quantification for four pseudobiological replicates, per strain, per experimental run. The data have been faceted by experimental run. Asterisks indicate the statistically significant difference of Student’s *t*-test between the indicated strain and wild-type Psa3 V-13, where p ≤.05 (*), p≤.01 (**), p≤.001 (***), and p>.05 (ns; not significant). These three experiments (biological replications) were separately conducted with three batches of independently grown plants.

**Figure S8:**
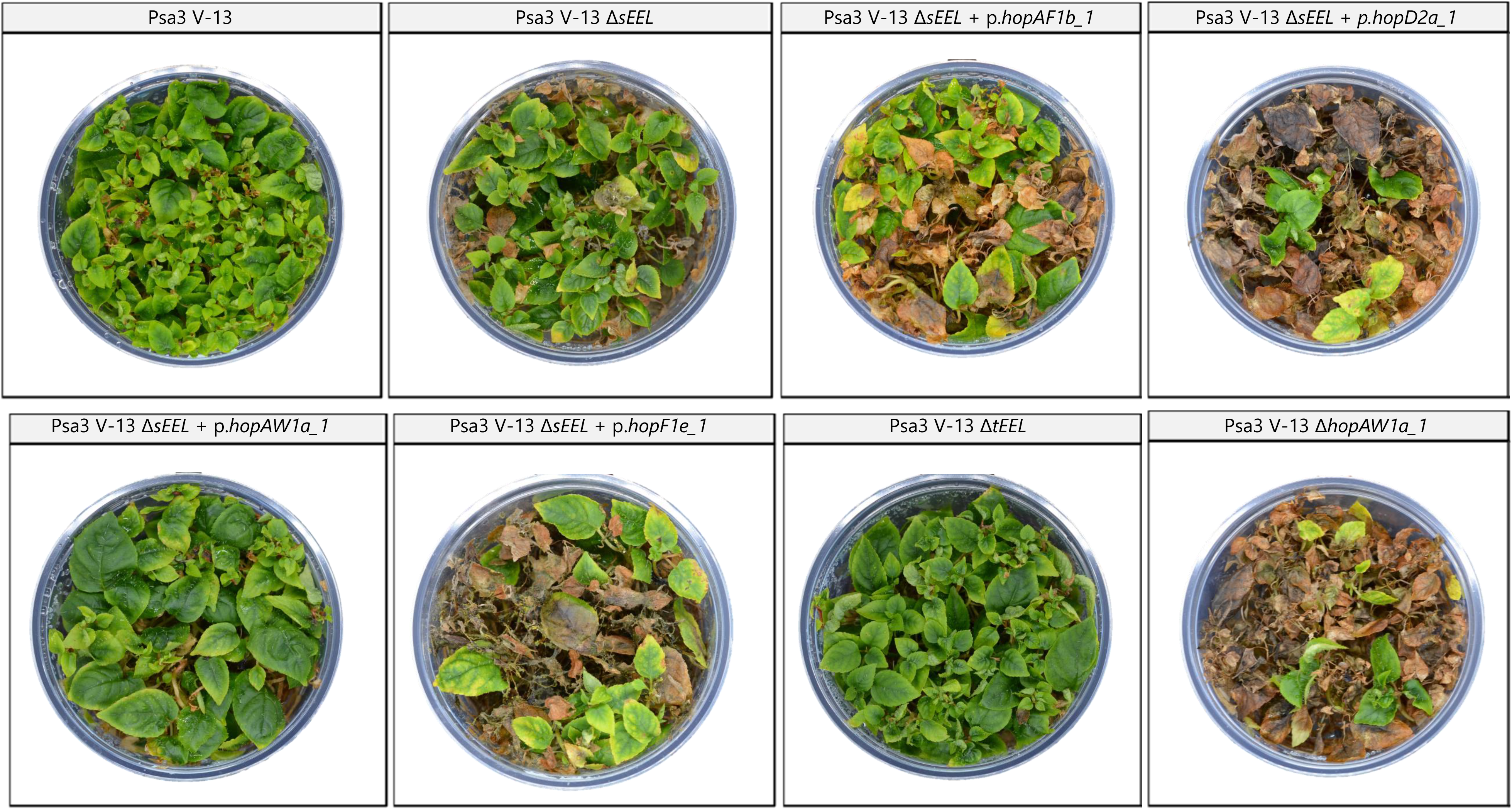
Symptom development of Psa3 V-13 *ΔsEEL* strains complemented with plasmids carrying individual sEEL effectors and Psa3 V-13 *ΔtEEL* and *ΔhopAW1a* strains in *Actinidia arguta*. *A. arguta* AA07_03 kiwifruit plantlets were flood-inoculated at approximately 10^7^ cfu/mL. Photographs of symptom development with representative pottles were taken at 50 days post-infection.

**Figure S9:**
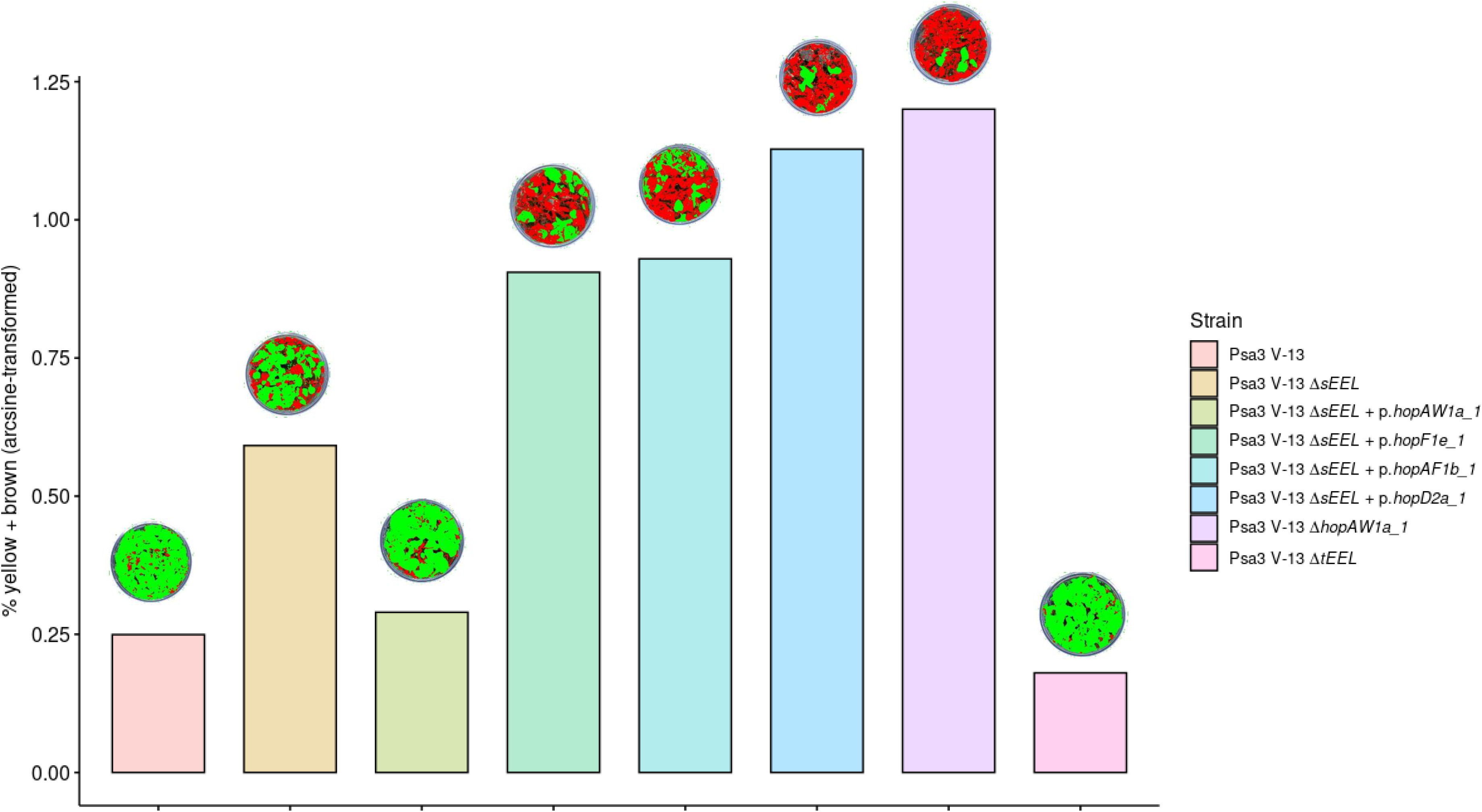
Quantification of symptom development of Psa3 V-13 *ΔsEEL* strains complemented with plasmids carrying individual sEEL effectors and Psa3 V-13 *ΔtEEL* and *ΔhopAW1a* strains in *Actinidia arguta.* A modified PIDIQ image-based analysis of leaf yellowing and browning, expressed as a normalized arcsine-transformed percentage for symptomology photographs taken at 50 days post-infection (Figure S7). Methodology adapted and modified from that of Laflamme et al. (2020).

**Figure S10:**
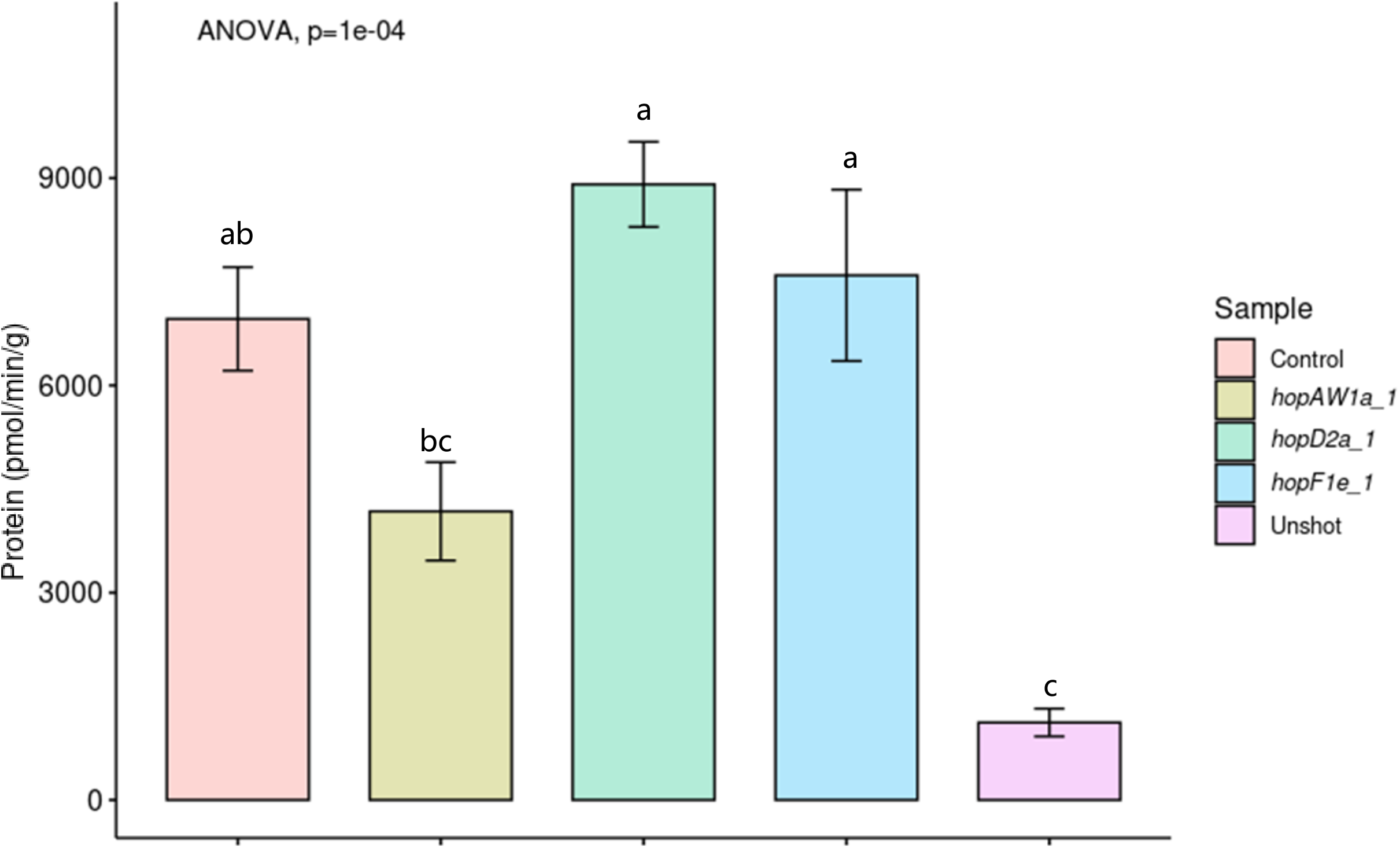
Biolistic transformation reporter eclipse assays demonstrate that HopAW1a is the sole sEEL effector triggering a hypersensitive response in *Actinidia arguta* AA07_03. *sEEL* effectors in cloned binary vector constructs tagged with GFP, or an empty vector (Control), were co-expressed with a β-glucuronidase (GUS) reporter construct using biolistic bombardment and priming in leaves from *A. arguta* AA07_03 plantlets (Jayaraman et al., 2021). The GUS activity was measured 48 hours after DNA bombardment. Error bars represent the standard errors of the means for three independent biological replicates with six technical replicates each (n=18). Un-infiltrated leaf tissue (Unshot) was used as a negative control. Tukey’s HSD indicates treatment groups which are significantly different at α ≤ 0.1 with different letters.

**Figure S11:**
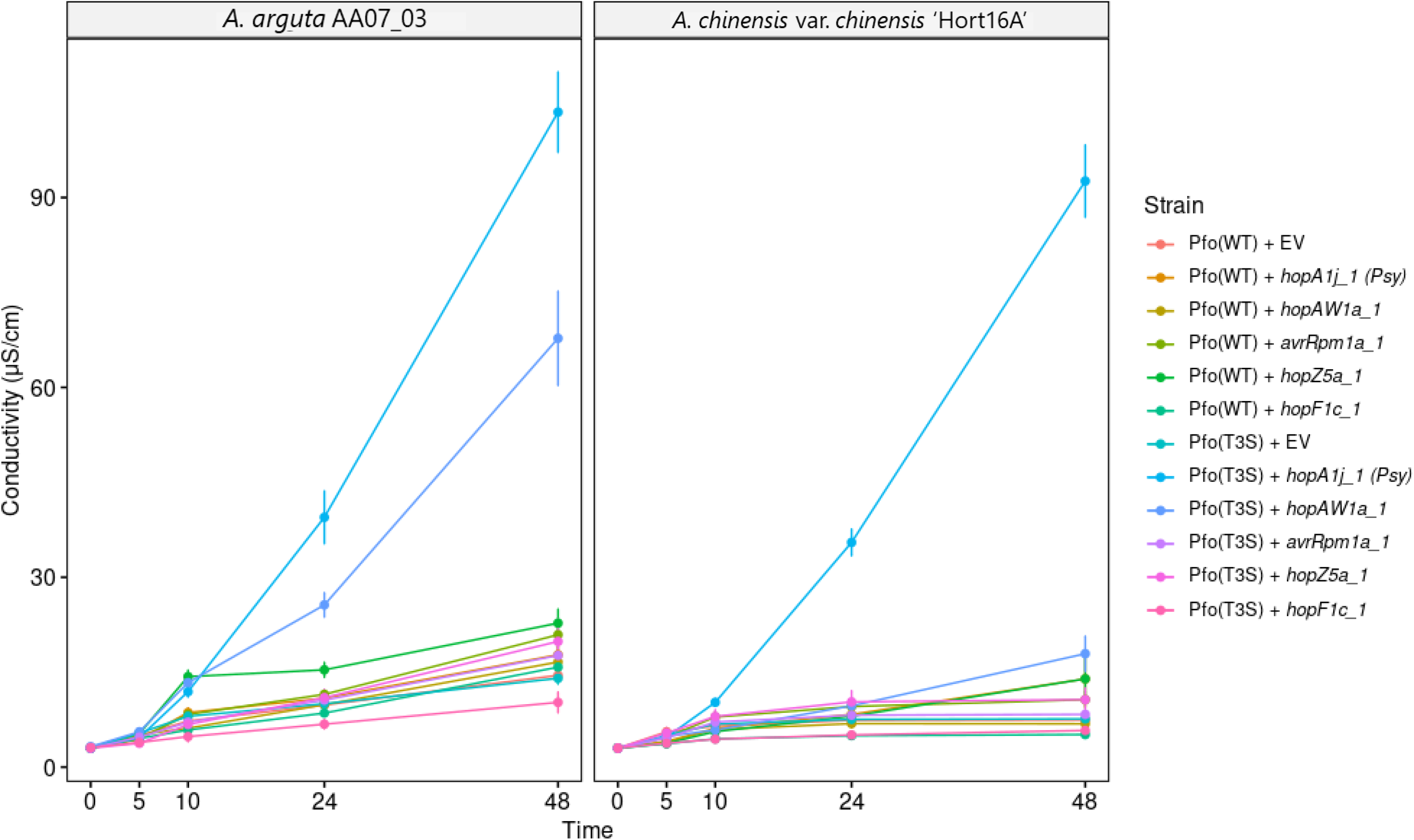
Measurement of hypersensitive response (HR) by ion leakage. Leaf discs from *A. arguta* AA07_03 and *A. chinensis* var. *chinensis* ‘Hort16A’ plantlets were vacuum-infiltrated with Psa3 inoculum at ∼5 x 10^8^ cfu/mL. Electrical conductivity due to HR-associated ion leakage was measured at selected time points over 48 hours. The ion leakage curves are faceted by plant species and stacked for three independent runs of this experiment. Error bars represent the standard errors of the means calculated from the five pseudobiological replicates per experiment (n=15).

**Figure S12:**
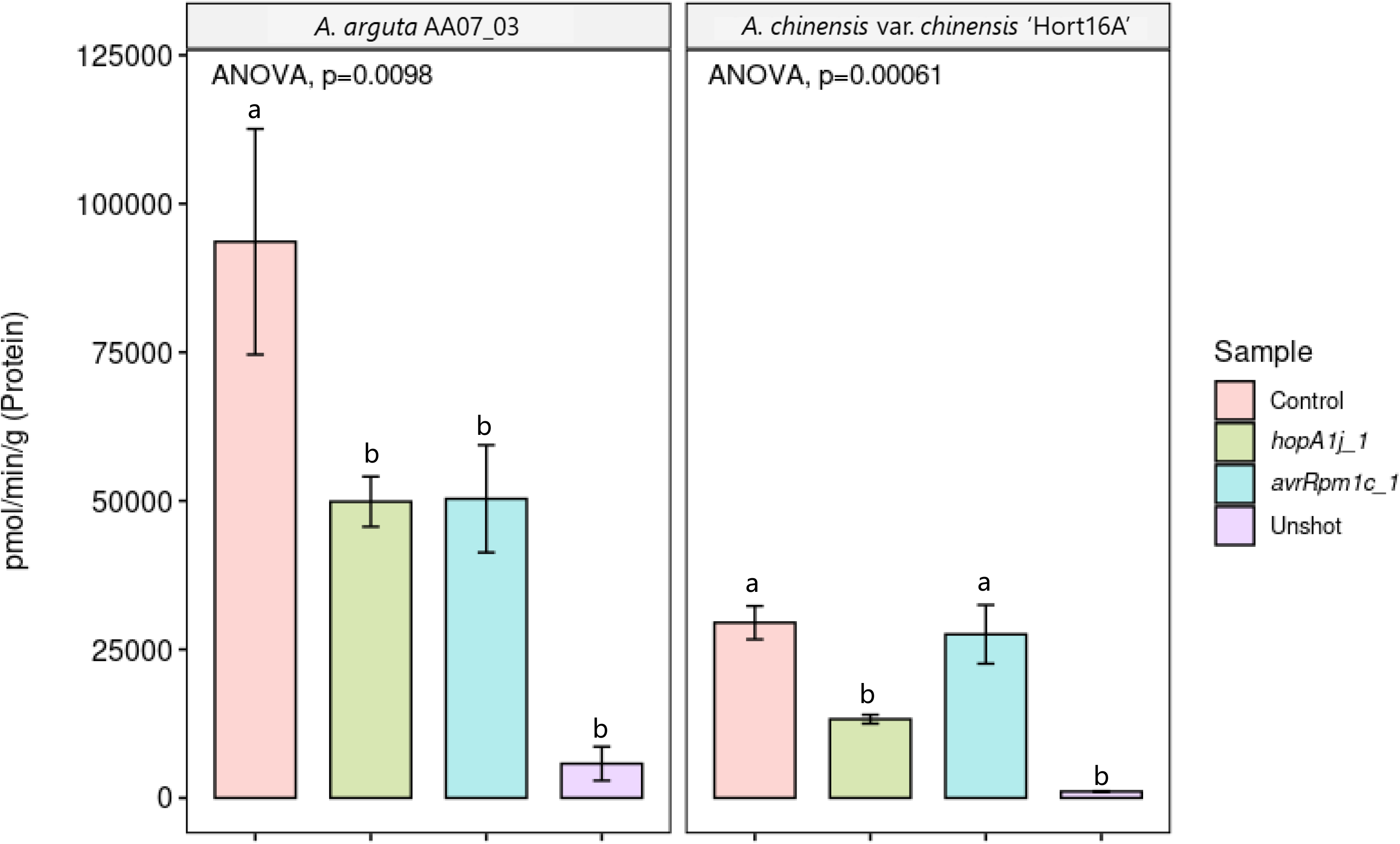
Biolistic transformation reporter eclipse assays demonstrate that AvrRpm1c from Psa2 K-28 triggers a host-specific immunity response in *Actinidia arguta* AA07_03. Effectors in cloned binary vector constructs tagged with Green Fluorescent Protein (GFP), or an empty vector (Control), were co-expressed with a β-glucuronidase (GUS) reporter construct using biolistic bombardment and priming in leaves from *A. arguta* AA07_03 plantlets (Jayaraman et al., 2021). The GUS activity was measured 48 hours after DNA bombardment. Error bars represent the standard errors of the means for three independent biological replicates with six technical replicates each (n=18). HopA1 from *Pseudomonas syringae* pv. syringae 61 was used as the positive control and un-infiltrated leaf tissue (Unshot) as the negative control. Tukey’s HSD indicates treatment groups which are significantly different at α ≤ 0.1 with different letters.

## Notes

### Competing Interest Statement

The authors have declared no competing interest.

